# Median Preoptic Astrocytes: Role in Sleep Regulation and Potential Mediators of Sex Differences

**DOI:** 10.1101/2025.08.27.672605

**Authors:** Chimdiya Onwukwe, Carissa A. Byrd, Shaun Viechweg, Destiny Black, Jessica Mong

## Abstract

One in three Americans suffer from chronic sleep disorders, and women are 40% more likely than men to experience sleep disorders. This disparity emerges at puberty and is strongly associated with fluctuations in the ovarian hormone, estrogen (E2), suggesting that E2 and biological sex are a risk factor for sleep disorders. Previous work in the lab has demonstrated that E2 suppresses sleep in female rats, including in sleep deprived rats whose homeostatic need for sleep is increased. However, the specific mechanism for E2 induced decrease in sleep remains unknown. Work in the lab suggests a role for adenosine in mediating E2’s sleep suppressive effects; E2 significantly increases Median Preoptic Nucleus (MnPO) extracellular adenosine and attenuates the action of specific agonists on the sleep promoting A2A-Receptor.

Astrocytes represent a major source of adenosine in the CNS and have been shown to influence neuronal activity and downstream behaviors. In this project, we tested the hypothesis that astrocytes mediate E2’s sleep suppressive effects. We used Gq-linked designer receptors exclusively activated by designer drugs (DREADDs) to evaluate the Gq pathway, which represents a core signaling mechanism in astrocyte activity. We found that, in female rats, activation of Gq signaling in astrocytes decreased sleep and inhibited homeostatic need for sleep. We further expressed the Pleckstrin Homology domain of PLC-like protein (p130PH), which has been shown to attenuate astrocyte activity and functions, in median preoptic nucleus (MnPO) astrocytes. We found that p130PH expression in MnPO astrocytes raised homeostatic sleep pressure to the same extent as 6 hours of sleep deprivation.

We further report that inhibiting astrocytic function did not prevent E2’s sleep suppressing effects suggesting that astrocytes may not play a role in estrogenic modulation of sleep. However, we did discover that MnPO astrocyte effects on sleep are sex-dependent. p130PH expression in MnPO astrocytes increased sleep and homeostatic sleep drive in female rats but showed a trend towards decreasing sleep and homeostatic sleep need in males. Further, while astrocyte effects on homeostatic sleep need are relegated to the dark phase in female rats, astrocytes appear to influence homeostatic sleep need in both the dark and light phase. To our knowledge, this is the first demonstration of a sex-based difference in astrocyte effects on sleep and homeostatic sleep pressure.

## Introduction

Sleep is an evolutionary conserved behavioral state crucial for maintaining bodily homeostasis. Homeostatic sleep drive is the body’s innate need for sleep and is typically heightened following periods of increased wakefulness. This sleep need is reflected by increased sleep duration following sleep loss, and enhanced synchronization of cortical activity as measured by electroencephalography (EEG). The cellular substrates underlying sleep homeostasis are not well understood. Despite this, sleep behavior and sleep homeostasis were previously believed to result from neuronal activity in key brain structures. However, recent studies suggest a role for non-neuronal cells, such as astrocytes, in sleep-wake behavior and homeostatic sleep drive.

Astrocytes are glial cells previously believed to constitute a purely structural component of the CNS. A growing body of work implicates astrocytes in controlling ionic balance in the extracellular space, modulating neuronal activity, and contributing to emergent behaviors including sleep. Manipulation of astrocyte G Protein Coupled Receptors in wake-promoting nuclei such as the basal forebrain and lateral hypothalamus increased wakefulness and decreased sleep. Meanwhile, disrupting astrocyte Ca^2+^ signaling and neuromodulator release decreased homeostatic sleep pressure. However, these studies have largely focused on astrocyte activity in areas of the brain that generate wake. There are few studies examining whether astrocytes in sleep-promoting nuclei regulate sleep and homeostatic sleep drive. The median preoptic nucleus (MnPO) regulates homeostatic sleep pressure and is one of two brain nuclei responsible for generating NREM sleep. MnPO GABAergic neurons promote sleep and signal sleep need, firing with increasing intensity during sleep deprivation and returning to baseline after recovery sleep^50^. However, it is not clear whether MnPO astrocytes also contribute to sleep behavior and homeostatic sleep need.

We investigated whether increasing MnPO astrocyte activity affects sleep-wake behavior and cortical markers of homeostatic sleep pressure. To do this, we expressed Gq associated Designer Receptors Exclusively Activated by Designer Drugs (Gq-DREADDs) in MnPO astrocytes. We chose this approach because endogenous astrocyte activity is largely driven by G Protein Coupled Receptors, and especially Gq-GPCRs expressed on the cell membrane. When activated, Gq-GPCRs increase the production of IP3 which liberates Ca^2+^ from the endoplasmic reticulum. Gq-DREADDs have largely been shown to mimic this signaling pathway.

To selectively express Gq-DREADD in MnPO astrocytes, we used 2 viral constructs: AAV5-GFAP-hM3DGq-mCherry and AAV5-gfaABC1D-hM3DGq-mCherry. AAV-GFAP-hM3DGq is primarily astrocyte-specific, especially when paired with optimized serotypes (e.g., AAV5 or AAV-DJ8) and where GFAP expression is robust ^131^. In addition, this virus has previously been used to target astrocytes in other hypothalamic nuclei^131^. However, minor off-target neuronal transduction have been reported for GFAP and other astrocytic promoters (i.e. GLAST, Aldh1l1, GLT-1) depending on delivery method or brain region^132–134^. To mitigate potential off target expression, we repeated the experiment using two different GFAP promoters.

We anticipate that decreasing MnPO astrocyte activity will induce sleep and increase homeostatic sleep pressure. We also clarify the effects of MnPO astrocytes on hemostatic sleep drive using a homeostatic sleep challenge in the form of 6 hour sleep-deprivation (SD). This method has previously been shown to produce saturating effects on markers of homeostatic sleep pressure including sleep duration following SD, NREM-SWA, and NREM delta power.

We first employed a chemogenetic approach utilizing Gi Designer Receptor Exclusively Activated by Designer Drugs (Gi-DREADDS) to decrease MnPO astrocyte activity. This construct is shown to decrease cellular activity in neurons, and early studies in astrocytes suggested that Gi DREADD may similarly decrease astrocyte activity. However, because recent results overwhelmingly suggest that Gi DREADDs increase astrocyte activity in several CNS nuclei, we also used a different secondary approach to decrease MnPO astrocyte activity. Specifically, we used Pleckstrin Homology domain of PLC-like protein (p130PH), a protein that has been shown to disrupt IP_3_ mediated Ca^2+^ signaling in astrocytes, as well as downstream behaviors^130,131,145^. We examine the effects of p130PH at baseline, and in sleep deprived animals.

Our previous work showed that E2’s effects on sleep are mediated by the MnPO, a hypothalamic nucleus that powerfully regulates sleep and homeostatic sleep drive. Infusing an estrogen receptor antagonist into the MnPO blocked systemic E2’s effects on sleep duration and homeostatic sleep need^42,58^. However, the cellular substrates underlying E2’s actions remain unknown.

A possible clue of the implicated cell type exists in E2’s effects on adenosinergic signaling in the MnPO. In addition to increasing wake, systemic E2 enhanced extracellular adenosine content in the MnPO. E2 also attenuated sleep-promoting Adenosine-2A (A2A) receptor signaling, creating a setting in which wake-promoting adenosine A1R may dominate^42^. Thus, it is possible that the source of this elevated adenosine may mediate E2’s effects on sleep duration and sleep homeostasis.

Astrocytes are a major source of adenosine in the CNS, releasing ATP that is rapidly converted to adenosine in the extracellular space. A growing body of evidence implicates astrocyte-derived adenosine in sleep homeostasis. Global astrocyte-specific knockdown of key enzymes in adenosine metabolism decreased homeostatic sleep drive^61^. Further, global impairment of gliotransmission disrupted sleep pressure in an adenosine A1-receptor dependent manner ^97^. Astrocytic G-protein coupled receptor pathways and astrocyte Ca^2+^ signaling also contribute to sleep-wake behavior^97,99^. Manipulation of astrocyte Gq signaling pathways in basal forebrain, lateral hypothalamus, and parabrachial nucleus increased wakefulness and decreased sleep^100–102^.

In addition, several studies show that astrocyte morphology, signaling and functions are extremely sensitive to modulation by estrogens^113,116,121,126,131,147,148^. A prenatal spike in E2 precipitates sex differences in astrocyte morphology in several CNS nuclei. Further, E2 has been shown to increase Ca^2+^ signaling in astrocytes in a sex-dependent manner.

In this study, we evaluated whether MnPO astrocytes mediate E2’s wake-promoting effects. We used p130PH which decreases astrocyte Ca^2+^ activity, to explore the contributions of astrocyte Ca^2+^ signaling to sleep. Given the sex differences in astrocyte morphology, we also explored whether there are sex differences in how MnPO astrocytes affect sleep and homeostatic sleep drive. Finally, we investigated how E2 affects the activity of cells in the MnPO.

## Results

### Activation of MnPO astrocyte Gq-DREADD (AAV5-GFAP-hM3DGq) in the dark phase promotes wake in female rodents

Previous work has demonstrated a role for astrocytes in wake-promoting nuclei in homeostatic sleep. In this set of experiments, we sought to determine the effect of activating MnPO astrocytes on sleep-wake behavior in female rats. We selectively expressed Gq-DREADD in MnPO astrocytes and performed IHC for S100B, an astrocytic protein, to confirm viral expression in astrocytes performed IHC for S100B, an astrocytic protein, to confirm viral expression in astrocytes (Fig 1A-D).

**Figure 1:**
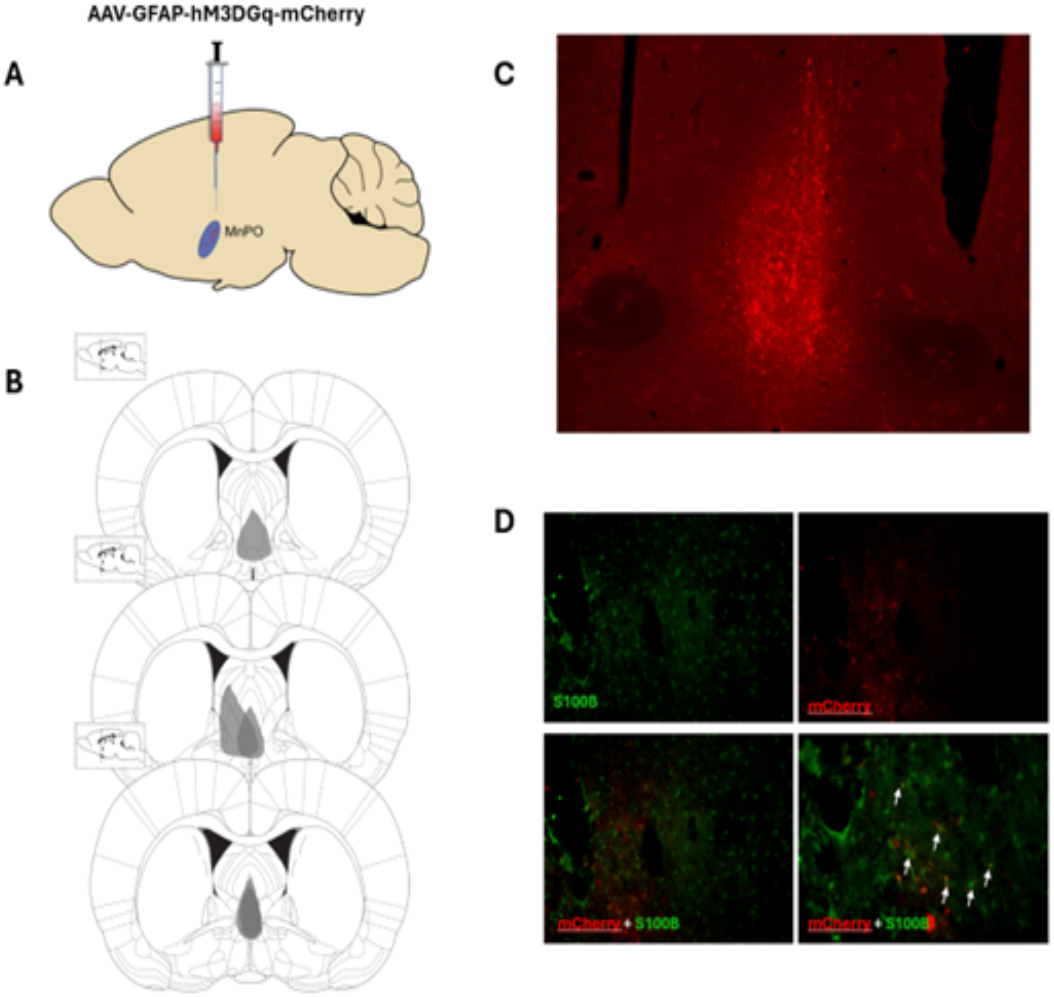
AAV-GFAP-hM3DGq-mCherry was used to express Gq-DREADD in MnPO astrocytes. (A) AAV-GFAP-hM3DGq was injected into the MnPO at Bregma. (B) Placement map of viral expression in experimental animals.(C) Representative image of viral expression in the MnPO (red=mCherry). (D) S100B-ir (green) was co-localized with viral expression (red).

In female rats, CNO injection at ZT0-1 significantly increased total wake time at the expense of NREM and REM sleep for 4 hours post-injection (Fig 2. B-D). This effect was largely driven by consolidation of wakefulness, as reflected by less frequent but longer wake bouts (Fig 2 E,H). Mixed effects analysis of sleep architecture also revealed a decrease in the number (Fig 2 F,G) and duration (Fig 2 I,J) of NREM and REM sleep bouts after CNO treatment. Notably, CNO injection at the start of the Dark Phase did not affect wake or NREM in the subsequent Light Phase, but increased REM, suggesting strong protection of REM sleep. These results suggest that MnPO astrocytes stabilize wake while decreasing sleep initiation and maintenance during the dark phase.

**Figure 1:**
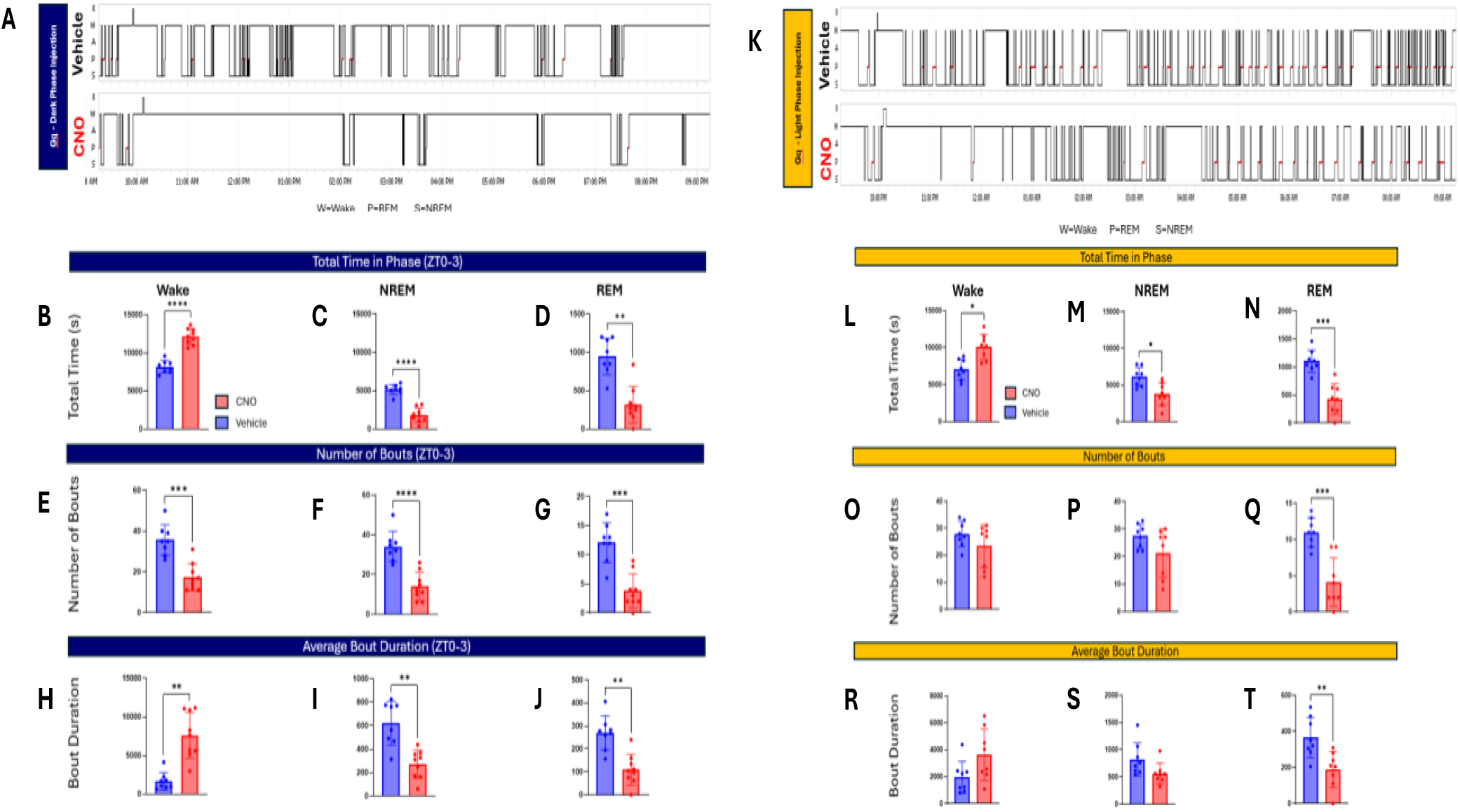
Chemogenetic activation of MnPO astrocytes during the Dark Phase and Light Phase increased wake and decreased sleep, in female rats. **(A)** Representative hypnogram of an animal treated with Saline Vehicle (top) or CNO (bottom) at the start of the dark phase. **(B-D)** Total time in wake, NREM and REM in the first 4 hours of the dark phase. CNO activation of MnPO astrocytes **(B)** increased total time in wake (paired t-test, t(7)=8.278, ****p<0.0001) and **(C)** decreased total time spent in NREM (paired t-test, t(7)=8.361, ****p<0.0001) and **(D)** REM (paired t-test, t(7)=5.119, **p=0.0014) in the first 4 hours of the dark phase. **(E-G)** Number of wake, NREM and REM bouts in the first 4 hours of the dark phase. CNO treatment decreased the number of **(E)** wake (paired t-test, t(7)=7.445, ****p=0.0009), **(F)** NREM (paired t-test, t(7)=8.842, ****p<0.0001) **(G)** and REM (paired t-test, t(7)=5.458, ***p<0.0001) bouts in the first 4 hours of the dark phase.**(H-J)** Average duration of wake, NREM and REM bouts in the first 4 hours of the dark phase. CNO treatment **(H)** increased the average duration of wake bouts (paired t-test, t(7)=4.551, **p=0.0026), and **(I)** decreased the average duration of NREM (paired t-test, t(7)=3.509, **p=0.0099) and **(J)** REM bouts. (paired t-test, t(7)=5.079, ****p=0.0014). **(K)** Representative hypnogram of an animal treated with Saline Vehicle (top) or CNO (bottom) at the start of the light phase. **(L-N)** Total time in wake, NREM and REM in first 4 hours of light phase. Chemogenetic activation of MnPO astrocytes **(L)** increased total time in wake (paired t-test, t(7)=3.321, *p=0.0128) and (**M)** decreased total time in NREM (paired t-test, t(7)=2.74, *p=0.0289) and **(N)** REM sleep (paired t-test, t(7)=6.046, ***p=0.0005) **(O-Q)** Number of wake, NREM and REM bouts in the first 4 hours of the light phase. Chemogenetic activation of MnPO astrocytes did not affect number of wake and NREM bouts but significantly decreased the number of REM bouts (paired t-test, t(7)=5.520, ***p=0.0009) **(R-T)** Average duration of wake, NREM and REM bouts in the first 4 hours of the light phase. Chemogenetic activation of MnPO astrocytes **(R)** showed a trend towards increasing average duration of wake bouts and **(S)** decreasing average NREM duration (paired t-test, t(7)=2.74, p=0.0981), and **(T)** significantly decreased average duration of REM (paired t-test, t(7)=5.135, **p=0.0013). Data are mean+/-SD for bar graphs and mean +/-SEM for hourly figures.

### Activation of MnPO astrocyte Gq-DREADD (AAV5-GFAPhM3DGq) in the light phase promotes wake in female rodents

Next, we investigated whether activation of MnPO astrocyte Gq-DREADD (AAV5-GFAP-hm3DGq) disrupts sleep during the light phase when the homeostatic need for sleep is high. Similar to our findings in the dark phase, CNO treatment in the light phase (ZT12-13) powerfully enhanced wake and reduced sleep (Fig 2 K). The first 4 hours of the light phase were particularly affected, with a significant increase in total wake time and marked reductions in total NREM and REM sleep (Fig 2 L-N). Unlike in the dark phase, CNO did not significantly affect the number or duration of wake (Fig 2 O,R) and NREM (Fig 2 P,S) bouts although there was a trend towards longer wake bouts and shorter NREM bouts. In contrast, the number and duration of REM bouts were significantly reduced (Fig 2 Q,T). Interestingly, in the subsequent Dark Phase, wake and REM were not affected but NREM was decreased.

Together, these findings demonstrate that MnPO astrocytes promote wake and suppress sleep in the dark and light phase. MnPO astrocyte’s ability to increase wake at the start of the light phase when sleep pressure is high suggests that astrocytes may influence the homeostatic need for sleep. In the next section, we evaluate the effect of activating MnPO astrocytes on cortical markers of sleep propensity.

### Activation of MnPO astrocytes Gq-DREADD (AA5-GFAP-hM3DGq) in the dark phase decreased cortical markers of homeostatic sleep pressure in female rats

To determine whether astrocytes influence cortical markers of sleep homeostasis, the PSD and NREM-SWA of CNO-treated animals were compared to Saline controls. Under physiological conditions, prolonged periods of wakefulness (>70 mins) increase markers of homeostatic sleep need including NREM delta power and NREM slow wave activity (NREM-SWA). While CNO injection at the start of the dark phase induced prolonged wakefulness, animals did not display the expected compensatory increases in markers of sleep need.

Comparisons of the PSD showed that CNO treatment reduced delta power (0.5-0.75Hz) in NREM bouts during the 12hr dark phase (Fig 3 A). Specifically, EEG spectral power was decreased in the 0.5-0.75Hz range across the entire dark phase. This effect may stem from significant reductions in NREM delta and theta power in the first 4 hours of the dark phase following CNO injection (Fig 3 B).

**Figure 3:**
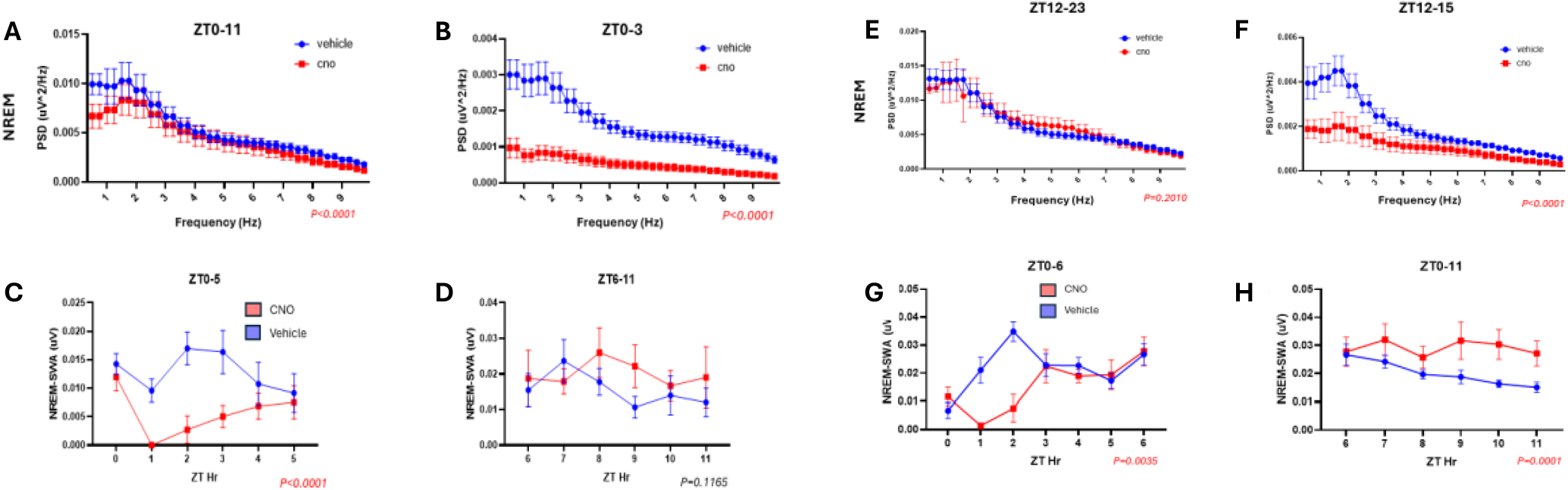
In female rats, activation of MnPO astrocytes during the Dark Phase and Light Phase decreased NREM delta power and NREM-SWA, a marker of sleep propensity, with compensatory rebounds later in the light phase. (A,B) NREM PSD in entire 12hr Dark Phase and in the first 4 hr of the Dark Phase. (**A**) CNO treatment decreased NREM delta power in the full 12 hour dark phase (Mixed Effects Analysis; main effect of CNO: F_(1, 228)_ =41.92, ****p<0.0001) (**B**) CNO treatment decreased NREM delta power in the first 4 hours of the dark phase (Mixed Effects Analysis; main effect of CNO: F_(1, 228)_ = 838.7, ****p<0.0001 **(C**,**D)** NREM-SWA in the first and last 6 hours of the Dark Phase. **(C)** CNO treatment decreased NREM-SWA in the first 6 hours of the dark phase (Mixed Effects Analysis; main effect of CNO: F_(1, 36)_=24.65, ****p<0.0001). **(D)** There was no compensatory rebound in NREM-SWA activity in the last 6 hours of the dark phase (Mixed Effects Analysis; main effect of CNO: F_(1, 36)_ = 2.587, p=0.1165). **(E**,**F)** NREM PSD in 12hr Light Phase and First 4hrs of the Light Phase. **(E)** CNO treatment did not significantly affect EEG spectral power in the 12-hour Light Phase. **(F)** However, in the first 4 hours of the Light Phase, CNO treatment significantly decreased EEG spectral power (RM 2-way ANOVA; main effect of CNO: F_(1, 228)_ = 312.8, ****p<0,0001), with post hoc tests revealing a significant decrease in NREM delta power specifically. Data are mean +/-SEM **(G**,**H)** NREM-SWA in the first and last 6 hours of the Light Phase. CNO treatment decreased NREM-SWA in the first 6 hours of the light phase (RM 2-way ANOVA; main effect of CNO: F(_1, 36)_ = 12.36, **p=0.0012) but **(H)** resulted in a compensatory increase in NREM-SWA in the last 6 hours of the light phase (RM 2 way ANOVA; main effect of CNO: F_(1, 36)_ = 10.76, **p=0.0023). Data are mean +/-SEM.

Analysis of hourly NREM-SWA also showed a significant decrease in NREM-SWA in the first 6 hours of the dark phase in CNO treated animals compared to Saline (Fig 3C). Notably, there was no compensatory rebound in NREM-SWA during the later parts of the dark phase (Fig 3D). We next examined other markers of sleep need, including inappropriate leakage of delta and theta power into wake, and increased delta power in REM sleep. While delta power decreased in REM sleep indicating low sleep propensity, low delta frequency ranges (0.5-0.75Hz) were increased in wake bouts. Thus, while our results overwhelmingly suggest that activation of Gq-DREADD in MnPO astrocyte reduced cortical markers of sleep pressure, some markers of sleep propensity were increased.

### Activation of MnPO astrocytes Gq-DREADD (AAV5-GFAP-hM3DGq) in the light phase decreased cortical markers of homeostatic sleep pressure in female rats

CNO activation of astrocyte Gq-DREADD at the start of the light phase also impacted cortical markers of sleep homeostasis. Comparisons of the PSD revealed that, in animals expressing Gq-DREADD, CNO treatment did not affect NREM delta power in the 12hr light phase but significantly decreased NREM delta power in the first 4 hours of the light phase (Fig 3 E,F).

MnPO astrocyte activation also decreased NREM-SWA in the first 6 hours of the light phase (Fig 3G). Interestingly, unlike in the dark phase, there was a rebound of NREM-SWA in the last 6 hours of the light phase (Fig 3H), suggesting strong protection of sleep pressure during the light phase. Despite this, other markers of sleep pressure were absent, including delta leakage into wake and REM.

### Activation of Gq-DREADD expressed using AAV5-gfaABC1D-hM3DGq-mCherry increased wake

We conducted a preliminary study expressing Gq-DREADD in MnPO astrocytes using AAV5-gfaABC1D-hM3DGq-mCherry. We verified Gq-DREADD expression in astrocytes using IHC for S100B, an astrocyte marker. We found that CNO treatment increased wake and decreased sleep in female rats. These findings support our observations above that that MnPO astrocytes stabilize wake and suppress sleep.

### CNO Injection Does not Affect Sleep in Animals not Expressing Gq-DREADD

To ensure that the results above are not due to off target effects of CNO, we injected female rodents not expressing Gq-DREADD with CNO (3mg/kg). We found that administering a higher dose of CNO (3mg/kg) than we used in our experiments did not affect sleep-wake behavior in animals not expressing DREADD virus (Fig 4). Further, just expressing Gq-DREADD in MnPO astrocytes of female rats did not affect sleep-wake architecture. Sleep-wake was not significantly different in animals expressing tDTomato and Gq-DREADD animals that were treated with Saline Vehicle (Fig 5). This suggests that sleep-wake effects we observed in the experiments above are not due to off-target CNO effects, or solely expressing Gq-DREADD in MnPO astrocytes with no CNO treatment.

**Figure 4:**
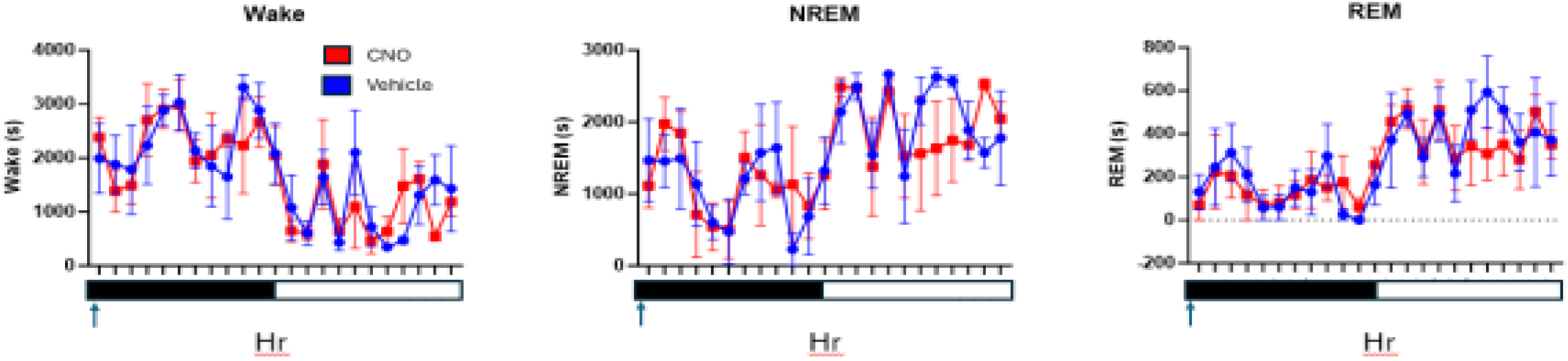
CNO treatment (3mg/kg) did not affect Sleep-Wake behavior in female rats not expressing Gq-DREADD. 24 hour time course for wake, NREM and REM after Vehicle or CNO treatment Data are mean+/-SD.

**Figure 5:**
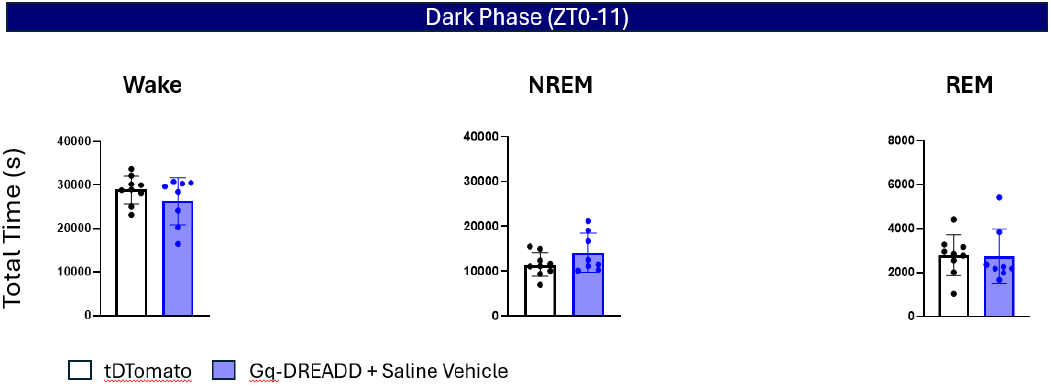
Just expressing Gq-DREADD in MnPO astrocytes did not significantly affect sleep-wake architecture. Total time in wake, NREM and REM for female rats expressing tDTomato or Gq-DREADD (with Saline Vehicle treatment) in MnPO astrocytes. Gq-DREADD expression in MnPO astrocytes did not significantly affect **(Left)** total wake time **(Middle)** total NREM time or **(Right)** total REM time compared to animals expressing tDTomato in MnPO astrocytes. Data are mean+/-SD.

### Activation of MnPO astrocyte Gi-DREADD in the dark phase promotes wake in female rodents

To determine the effects of decreasing astrocyte activity on Sleep, we expressed Gi-DREADD, an inhibitory GPCR in MnPO astrocytes and used polysomnographic recordings to collect EEG/EMG signals. We performed IHC for GFAP protein to confirm viral expression in the MnPO astrocytes (Fig 6) Surprisingly, CNO activation of astrocyte Gi-DREADD produced similar effects as Gq-DREADD, promoting wake and reducing NREM and REM sleep in the dark phase (Fig 7 A,B).

**Figure 6:**
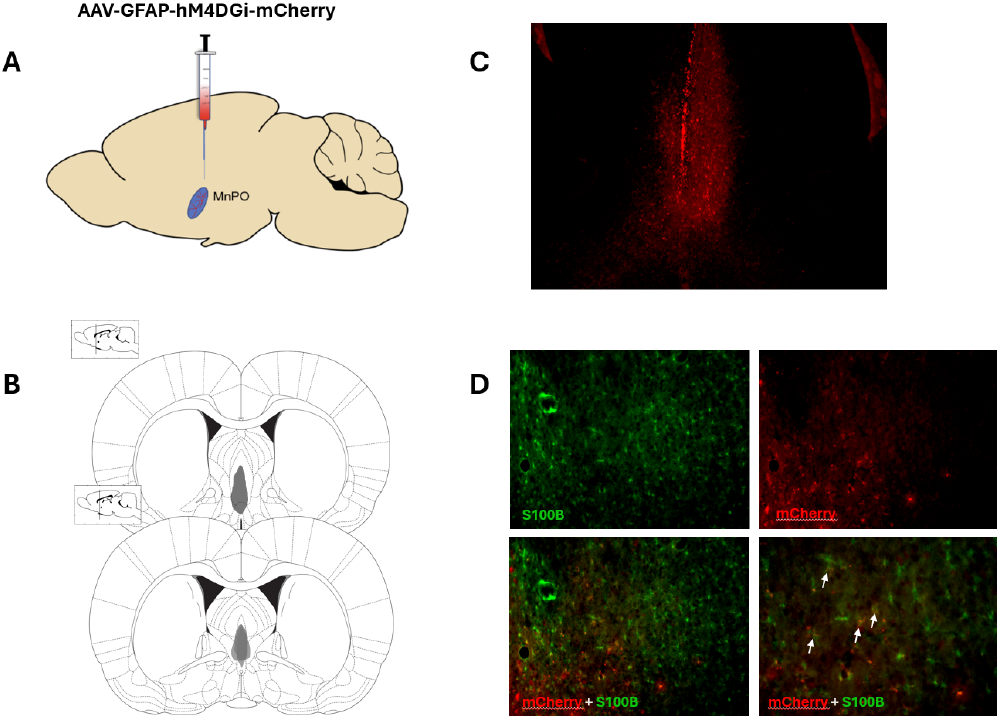
AAV-GFAP-hM4DGi-mCherry was used to express Gi-DREADD in MnPO astrocytes. (A) AAV-GFAP-hM4DGi was injected into the MnPO at Bregma. (B) Placement map of viral expression in experimental animals. (C) Representative image of viral expression in the MnPO (red=mCherry). (D) S100B-ir (green) was co-localized with viral expression (red).

**Figure1.7:**
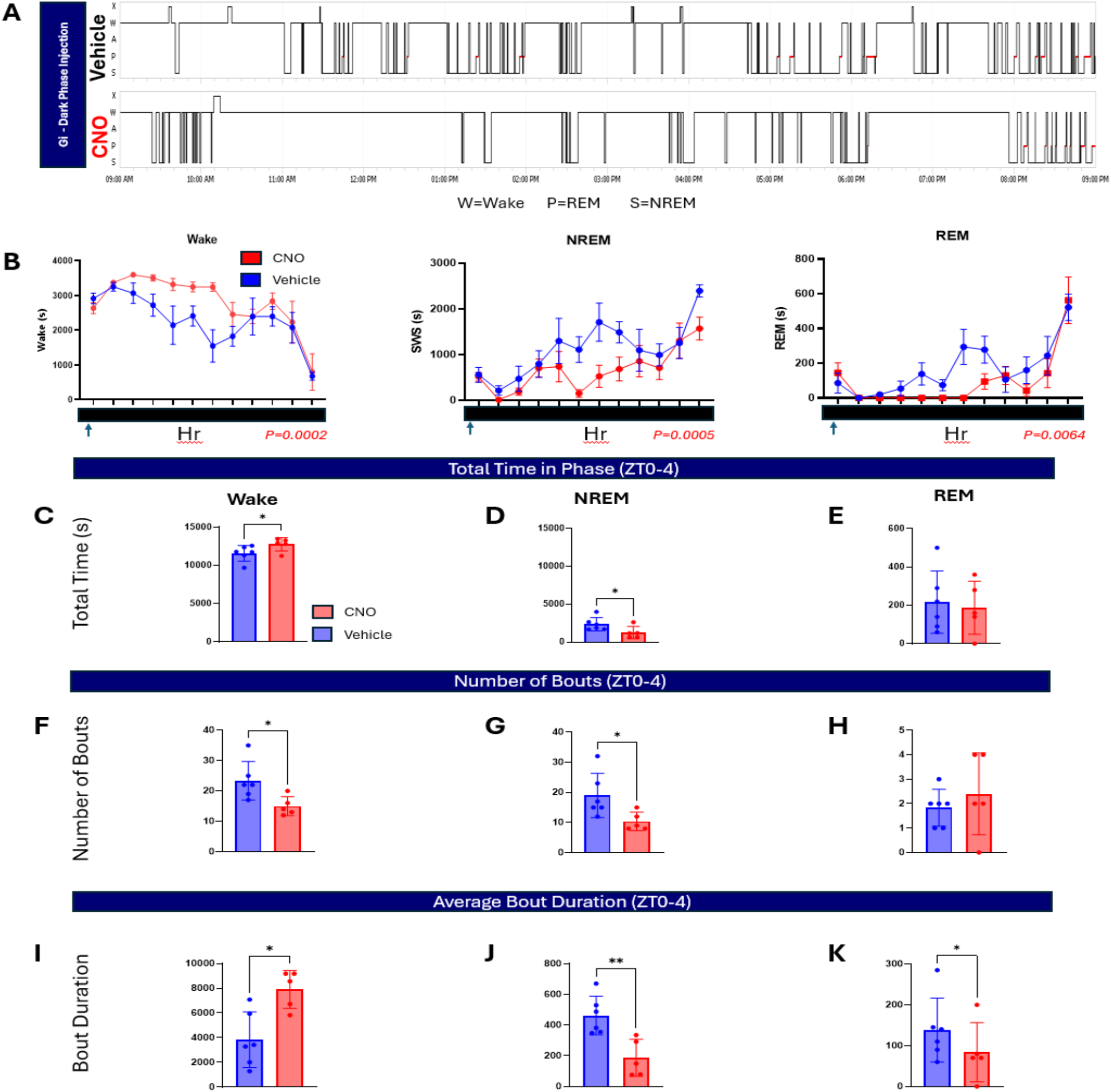
CNO activation of Gi-DREADD during the Light Phase increased wake and decreased Sleep in female rats. **(A)** Representative hypnogram of an animal treated with Saline Vehicle (top) or CNO (bottom) at the start of the dark phase. **(B)** Hourly sleep-wake states for Dark Phase (12 hours). Chemogenetic activation of Gi-DREADD in female rats (n=5) increased wake **(Left)** (Mixed Effect Model; main effect of CNO: F_(1, 36)_=17.09, ***p=0.0002) and decreased NREM **(middle)** (Mixed Effect Model; main effect of CNO: F_(1, 36)_ = 14.63, ***p=0.0005), and REM (Right) (Mixed Effect Model; main effect of CNO: F_(1, 36)_ = 8.398, **p=0.0064) sleep in the 12-hour time course. **(C-E)** Total time in wake, NREM and REM in the first 4 hours of the dark phase. CNO activation of Gi-DREADD **(C)** increased total time in wake (paired t-test, t(4)=3.430, *p=0.0265) and **(D)** decreased total time spent in NREM (paired t-test, t(4)=3.978, *p=0.0164) but not **(E)** REM. **(F-H)** Number of wake, NREM and REM bouts in the first 4 hours of the dark phase. CNO treatment decreased the number of **(F)** wake (paired t-test, t(4)=4.182, *p=0.0139), **(G)** NREM (paired t-test, t(4)=3.213, *p=0.0325) **(H)** but not REM. **(I-K)** Average duration of wake, NREM and REM bouts in the first 4 hours of the dark phase. CNO treatment **(I)** increased the average duration of wake bouts (paired t-test, t(4)=3.479, *p=0.0254), and **(J)** decreased the average duration of NREM (paired t-test, t(4)=5.465, **p=0.0055) and **(K)** REM bouts. (paired t-test, t(4)=3.315, *p=0.0295). *Data are mean+/-SD for bar graphs, and mean +/-SEM for hourly figures*.

These effects were driven by a consolidation of wake, with longer wake bouts on average, and less frequent wake, NREM and REM bouts (Fig 7 D, E).

### p130PH expression in MnPO astrocytes did not block E2’s wake-promoting effects

E2 treatment and activation of MnPO astrocytes produce similar effects on sleep in female rats, prompting us to investigate whether astrocytes mediate E2’s wake-promoting effects. To test this hypothesis, we used p130PH to reduce the activity of MnPO astrocytes in OVXed female rats, and tDTomato as a control (Fig 8).

**Figure 8:**
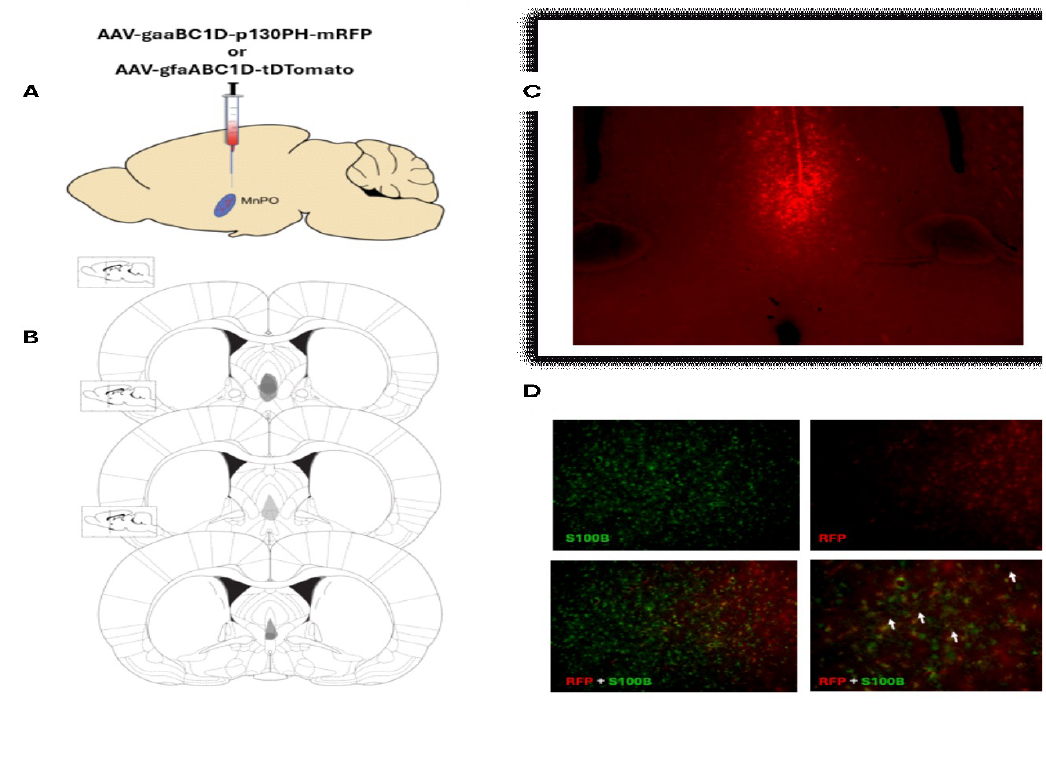
AAV-gfaaBC1D-p130PH-mRFP was used to express p130PH in MnPO astrocytes. **(A)** AAV-gfaaABC1D-p130PH-mRFP or AAV-gfaaBC1D-tDTomato was injected into the MnPO at Bregma. **(B)** Placement map of viral expression in experimental animals. **(C)** Representative image of viral expression in the MnPO (red=mRFP). **(D)** S100B-ir (green) was co-localized with viral expression (mRFP, red).

In agreement with our previous findings, E2 treatment significantly increased total wake time and decreased total NREM and REM sleep time in Control animals. In animals expressing p130PH, E2 administration similarly increased wake and decreased NREM and REM sleep (Fig 9). Thus, attenuating MnPO astrocyte activity with p130PH did not block E2’s wake-promoting effects.

**Figure 9:**
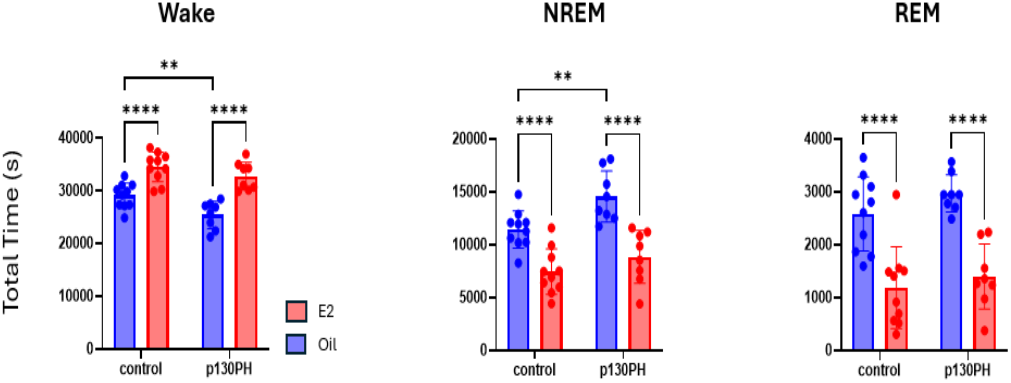
p130PH expression in MnPO astrocytes did not block E2’s wake-promoting effects in female rats. Total time in wake (left), NREM (middle) and REM (right). **(Left)** E2 increased wake in both p130PH expressing animals and Control animals. **(Middle)** E2 decreased NREM sleep in p130PH and Control animals. **(Right)** E2 decreased REM sleep in p130PH and Control animals. Data are mean+/-SD.

### Sex differences in astrocyte effects on sleep

Astrocyte morphology is significantly affected by a prenatal spike in E2 that male, but not female rats experience. These sex-based differences in astrocyte morphology have been observed in several hypothalamic nuclei and persist into adulthood. However, it is not known if there are sex differences in astrocyte function particularly in sleep. In this experiment, we examined whether there are sex differences in astrocyte effects on sleep behavior and homeostatic need for sleep. To do this, we expressed p130PH in MnPO astrocytes of male rats, and confirmed colocalization of virus in astrocytes.

We found that p130PH expression in MnPO astrocytes impacted sleep and homeostatic sleep drive differently in males and females. Specifically, p130PH expression increased sleep in female rats, but showed a trend towards decreasing sleep in male rats (Fig 10 A-C). Comparisons of the sleep-wake architecture revealed that p130PH did not affect the number of wake, NREM or REM

**Figure 10:**
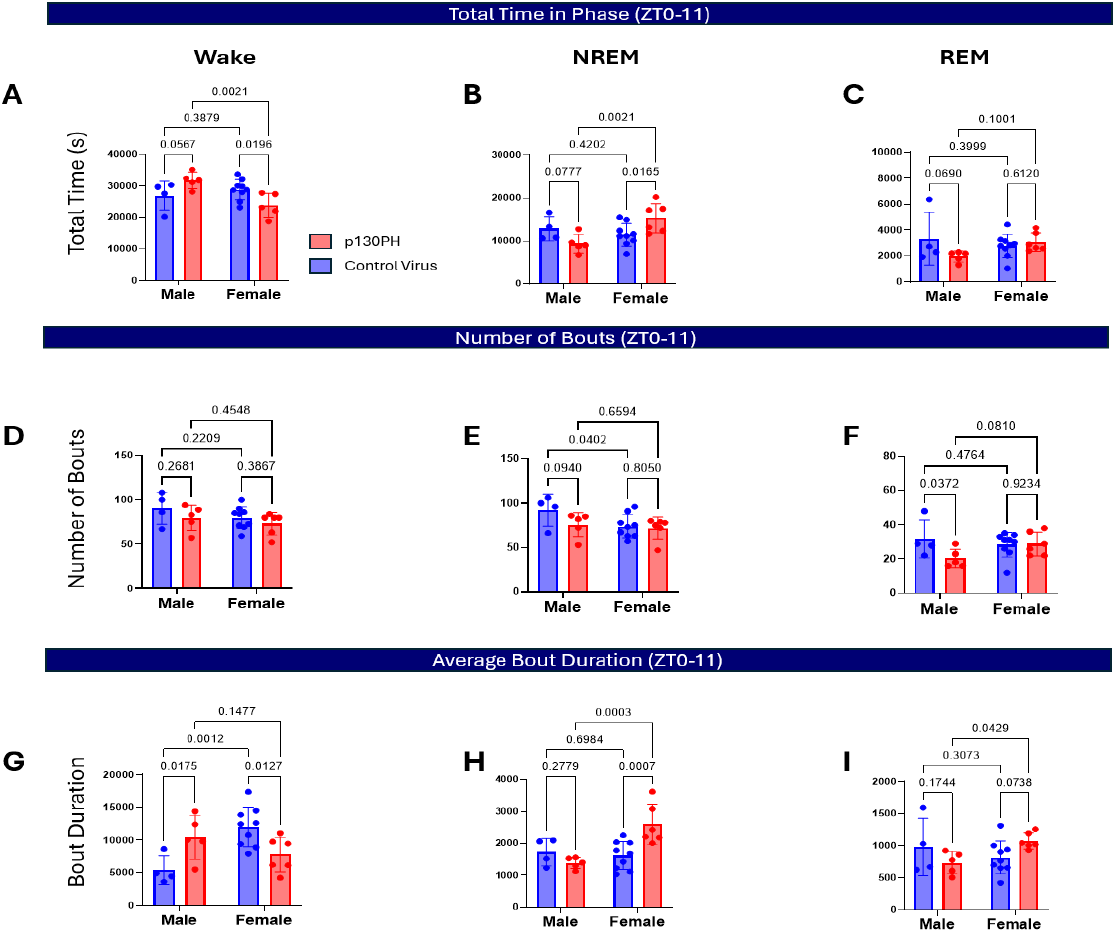
p130PH expression increased sleep in female rats, but showed a trend towards decreasing sleep in male rats. **(A-C)** Total time in wake, NREM and REM in dark phase for male and female rats. When animals expressing p130PH in MnPO astrocytes were compared to control rats: **(A)** total wake time was decreased in female rats, but showed a trend towards increasing in male rats **(B)** total NREM time was increased in female rats, but showed a trend towards decreasing in male rats **(C)** total REM time is not changed in female rats, but showed a trend towards decreasing in male rats. **(D-F)** Number of bouts in wake, NREM and REM in dark phase for male and female rats. **(D)** p130PH expression in MnPO astrocytes did not affect number of wake bouts in male or female rats. **(E)** p130PH in astrocytes did not affect number of NREM bouts in female rats but showed a trend to-wards decreasing number of NREM bouts in male rats. **(F)** p130PH expression does not affect number of REM bouts in female rats but showed a trend towards decreasing number of REM bouts in male rats. **(G-I)** Average duration of wake, NREM and REM bouts. In animals expressing p130PH: **(G)** Average duration of wake bouts was decreased in females but increased in males. **(H)** Average duration of NREM bouts was increased in females but not affected in males. Similary, **(I)** average duration of REM sleep bouts was increased in females but not affected in males.

We found that p130PH expression in MnPO astrocytes impacted sleep and homeostatic sleep drive differently in males and females. Specifically, p130PH expression increased sleep in female rats, but showed a trend towards decreasing sleep in male rats (Fig 10 A-C). Comparisons of the sleep-wake architecture revealed that p130PH did not affect the number of wake, NREM or REM bouts in female rats. In contrast, p130PH decreased the number of REM bouts in male rats and showed a trend towards decreasing the number of NREM bouts as well (Fig 10 D-F). Further, p130PH decreased the average length of wake bouts in female rats but increased the length of wake bouts in males (Fig 10 G).

P130PH expression also affected cortical markers of sleep pressure differently in male and female rats. In female rats, p130PH *increased* NREM-SWA, in the dark but not the light phase (Fig 11 A). In contrast, in male rats, p130PH *decreased* NREM-SWA in the dark and light phase (Fig 11 B). This suggests that p130PH expression in MnPO astrocytes increases sleep pressure in female rats but decreases sleep pressure in male rats.

**Figure 11:**
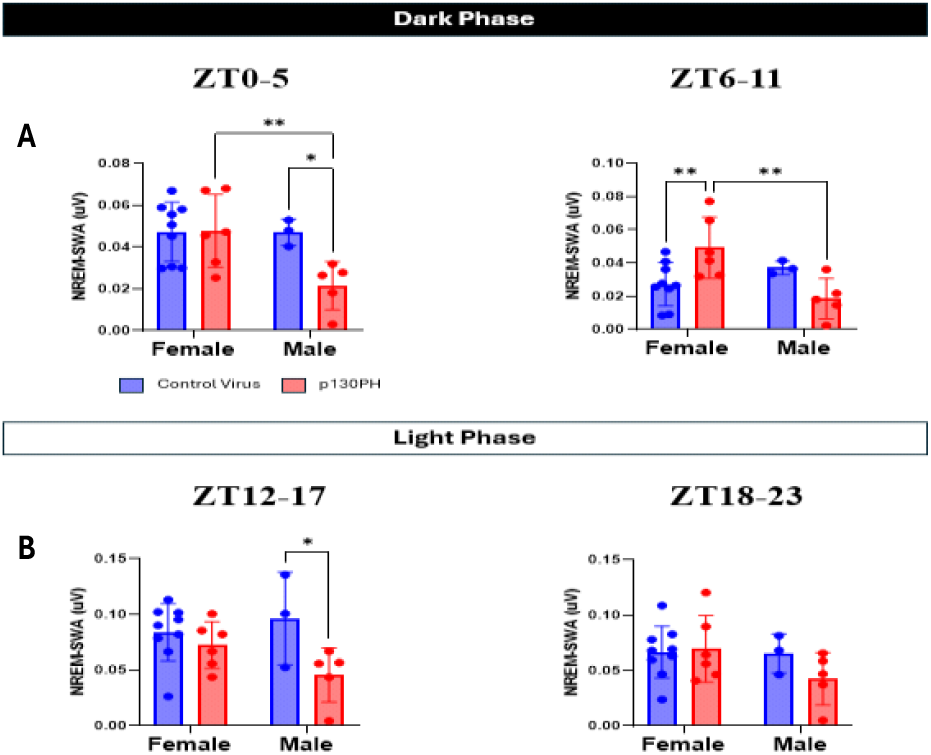
P130PH expression in MnPO astrocytes increased NREM-SWA in female rats during the dark phase, but decreased NREM-SWA in male rats in both light and dark phase. **(A)** 6-hour totals for NREM-SWA in the first and last half of dark phase for males and females. **(Left)** In the first 6 hours of the dark phase, p130PH expression did not affect NREM-SWA in female rats but decreased NREM-SWA in male rats. **(Right)** In the last 6 hours of the dark phase, p130PH increased NREM-SWA in females but decreased NREM-SWA in males. **(B)** 6-hours totals for NREM-SWA in the first and last half of light phase for males and females. **(Left)** In the first 6 hours of the light phase, p130PH expression does not affect NREM-SWA in female rats, but **(Right)** decreased NREM-SWA in males.

To better understand the sex-dependent effects of p130PH on homeostatic sleep need, we sleep deprived male rats. In p130PH males, SD increased recovery sleep and reduced wake in the dark, but not the light phase. This contrasts with our findings in p130PH females, where SD did not affect recovery sleep in the dark phase, but increased recovery sleep and reduced wake in the light phase (Fig 12).

**Figure 12:**
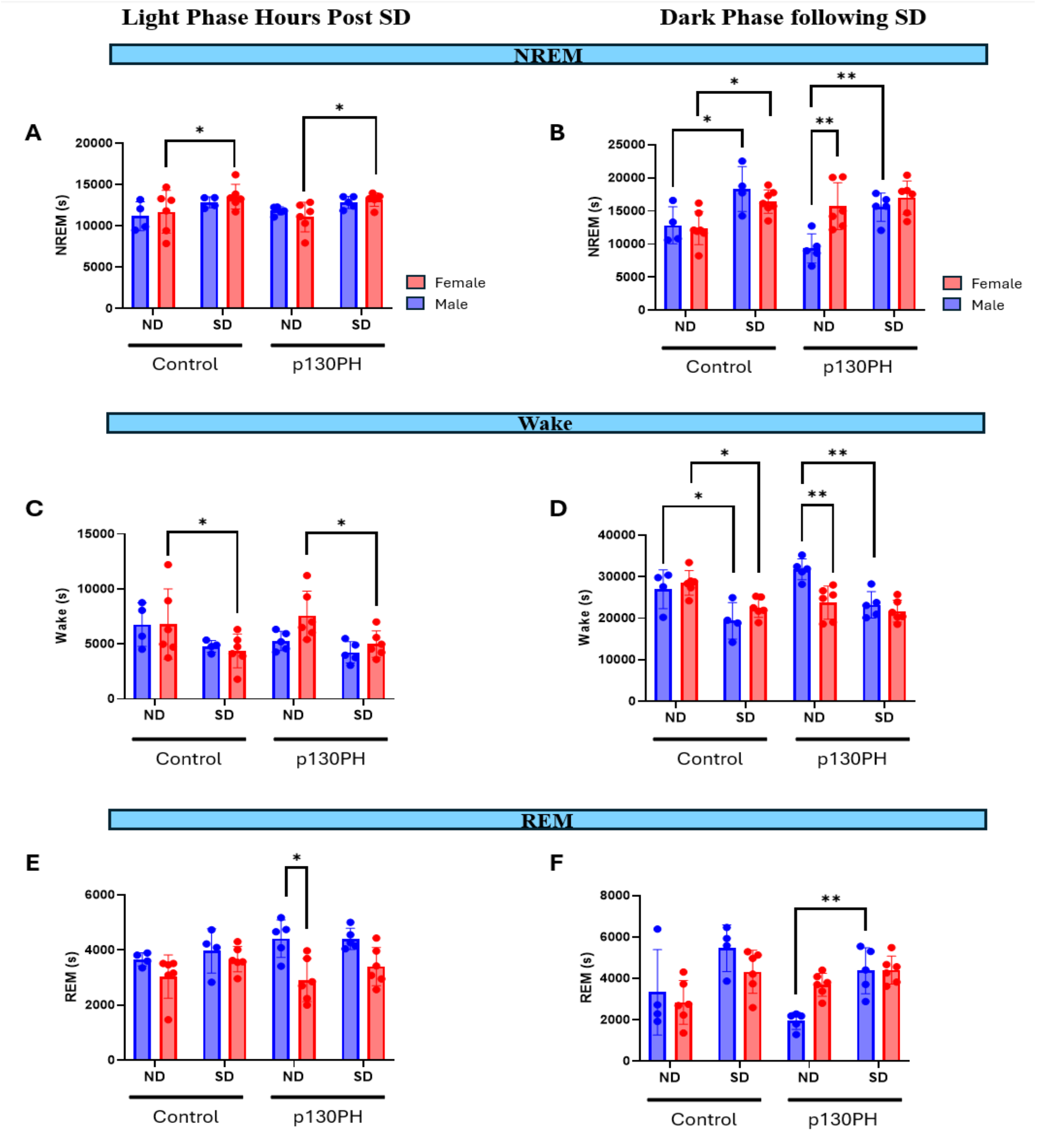
To explore whether there is a sex difference in the response to sleep deprivation (SD) between males and females, we ran a 3-way ANOVA with the factors being sex, virus (p130PH), and SD. (A) NREM sleep in the first 6 hours of the light phase following SD. There was a main effect of SD on NREM in the first 6 hours of light phase after SD (F_(1, 17)_ = 37.71, p<0.0001). Tukey post hoc test showed that SD significantly increased NREM in both control (p=0.0301) and p130PH female rats (p=0.0107). **(B) NREM sleep in the subsequent dark phase following light phase SD**. In the dark phase following SD, there was a main effect of SD on NREM (F_(1, 18)_ = 52.59, p<0.0001), and interactions between sex and p130PH (F_(1, 18)_ = 6.746, p=0.0182) as well as sex and SD (F_(1, 18)_ = 7.499, p=0.0135). Tukey post hoc test revealed that SD increased NREM in control rats regardless of sex (Male: p=0.0103; Female: p=0.0177). SD also increased NREM in male p130PH rats (p=0.0013), but not in female p130PH. Finally, there was a significant difference in NREM of male and female p130PH rats (p=0.0061). **(C) Wake in the first 6 hours of the light phase following SD**. There was main effect of SD on wake in the first 6 hours of the light phase after SD (F_(1, 17)_ = 29.52, p<0.0001). Tukey post test showed that SD significantly decreased wake in female control (p=0.0378) and p130PH rats (p=0.0249).**(D) Wake in the subsequent dark phase following light phase SD**. In the dark phase following SD, there was a main effect of SD on wake (F_(1, 17)_ = 57.34, p<0.0001), and interactions between sex and p130PH (F_(1, 17)_ = 8.257, p=0.0105) as well as sex and SD (F_(1, 17)_ = 6.533, p=0.0205). Tukey post hoc test revealed that SD increased NREM in control rats regardless of sex (Male: p=0.0105; Female: p=0.0154). SD also increased NREM in male p130PH rats (p=0.0012), but not in female p130PH. Finally, there was a significant difference in NREM of male and female rats expressing p130PH (p=0.0004). **(E) REM in the first 6 hours of the light phase following SD**. There were main effects of sex (F_(1, 17)_ = 12.55, p=0.0025) as well as SD (F_(1, 17)_ = 5.449, p=0.0321). Tukey post hoc analysis showed that REM was significantly different in ND p130PH males and females (p=0.0105). **(F) REM in the subsequent dark phase following light phase SD**. There was a main effect of SD (F _(1, 17)_ = 33.71, p<0.0001) and an interaction between sex and p130PH (F_(1, 17)_ = 5.653, p=0.0294). Post hoc analysis showed that SD significantly reduced REM in p130PH males (p=0.0127). Data are mean+/-SD.

## Discussion

Our previous work showed that E2 regulates sleep duration and homeostatic sleep drive. In this work, we investigated whether astrocytes mediate E2’s inhibitory effects on sleep. The current set of findings demonstrate that MnPO astrocytes affect sleep behavior and sleep homeostasis in a sex-dependent manner, but do not mediate E2’s effects on sleep-wake. Specifically, we found that Gq activation in MnPO astrocytes decreased sleep and sleep pressure in female rats. We also expressed p130PH, an IP3 chelator that has been shown to reduce astrocyte Ca^2+^ signaling, in MnPO astrocytes. p130PH expression in MnPO astrocytes increased sleep and sleep pressure in female rats but showed a trend towards decreasing sleep and sleep pressure in males. However, p130PH expression in MnPO astrocytes did not prevent E2’s sleep-suppressing effects in female rats. Thus, while E2 and astrocytes each reduce sleep in female rodents, E2’s sleep effects are not mediated by astrocytes.

### Activation of Gq-signaling in astrocytes increased wake in female rats

Astrocytes were previously thought to represent a purely structural component of the CNS. An emerging field of study indicates that astrocytes are involved in sleep behavior^100,102,107,108,111^. Chemogenetic activation of astrocyte Gq in wake-active nuclei, such as the BF, PB and LH, has been shown to promote wakefulness and inhibit sleep^100–102^. However, the role of astrocytes in sleep-promoting nuclei in sleep regulation remains understudied. Further, to our knowledge, there are no studies specifically evaluating astrocyte Gq effects on sleep in female rats.

In this work, we explored the role of astrocytes in the MnPO, one of two nuclei that has been implicated in sleep initiation. Our findings show that, in female rats, Gq activation in MnPO astrocytes increased wake without the compensatory increase in cortical markers of homeostatic sleep need that typically accompanies periods of prolonged wakefulness. In fact, Gq activation in MnPO astrocytes decreased NREM-SWA, a marker of homeostatic sleep pressure and sleep intensity.

One potential explanation for our findings is that Gq activation in MnPO astrocytes enhances release of gliotransmitters such as D-serine, adenosine, prostaglandin E2 (PGE2) and glutamate. These gliotransmitters may then suppress the activity of sleep-promoting MnPO neurons. Supporting this hypothesis, infusions of adenosine or A1R agonists in preoptic nuclei powerfully promotes wake^59,141^. In addition, gliotransmitters like PGE2 have been shown to suppress neuronal activity and sleep when infused into the preoptic area ^142^. Alternatively, Gq activation in astrocytes may enhance astrocytic clearance of excitatory transmitters and K^+^ ions from the extracellular space, hyperpolarizing sleep-promoting MnPO neurons. Astrocytes may and K^+^ ions from the extracellular space, hyperpolarizing sleep-promoting MnPO neurons. Astrocytes may

One potential explanation for our findings is that Gq activation in MnPO astrocytes enhances release of gliotransmitters such as D-serine, adenosine, prostaglandin E2 (PGE2) and glutamate. These gliotransmitters may then suppress the activity of sleep-promoting MnPO neurons. Supporting this hypothesis, infusions of adenosine or A1R agonists in preoptic nuclei powerfully promotes wake^59,141^. In addition, gliotransmitters like PGE2 have been shown to suppress neuronal activity and sleep when infused into the preoptic area ^142^. Alternatively, Gq activation in astrocytes may enhance astrocytic clearance of excitatory transmitters and K^+^ ions from the extracellular space, hyperpolarizing sleep-promoting MnPO neurons. Astrocytes may specifically inhibit MnPO GABAergic neurons which have previously been shown to promote sleep and homeostatic sleep pressure via inhibitory projections to wake-promoting nuclei such as the BF, LC and TMN.

Our data align with emerging studies demonstrating a role for astrocytes in homeostatic sleep. While most studies only report an effect of Gq activation in astrocytes on sleep-wake duration, a recent study by Ingiosi et al showed that Gq activation in astrocytes in the wake-active BF increased wake without compensatory increases in markers of sleep need^100,102,106,107^. Together with these findings, our observations in astrocytes demonstrate that sleep is not solely a product of neuronal activity as previously thought and highlights a need for further investigation into how astrocytes may fit into our current understanding of sleep-wake neurocircuitry.

### P130PH expression in MnPO astrocytes increases sleep in female rats

Several groups report an association between astrocyte Ca^2+^ signaling and the sleep-wake cycle. For example, in the hippocampus, cerebellum and cortex, astrocyte Ca^2+^ is high during wake, and low during sleep^103^. Moreover, our data suggests that astrocyte Gq activation in the MnPO increased wake. Because the Gq pathway has repeatedly been shown to increase astrocyte Ca^2+^ levels, it is likely that this effect is mediated by astrocyte Ca^2+^ signaling. To clarify the contributions of astrocyte Ca^2+^ to sleep, we expressed p130PH, which disrupts IP_3_-Ca^2+^, in MnPO astrocytes.

p130PH expression in MnPO astrocytes increased sleep in the ***<u>dark</u>***, but not the light phase. Strikingly, these effects were most prominent in the last 6 hours of the dark phase. This temporal profile aligns with previous studies indicating that astrocytes are most active during the dark phase and may suggest involvement of the gliotransmitter adenosine which peaks in the latter half of the dark phase^96,109,153^. Specifically, p130PH expression may attenuate astrocytic release of adenosine, thereby reducing A1R mediated inhibition of sleep-promoting neurons.

Our findings do not align with other studies examining the effects of attenuating astrocyte Ca^2+^ on sleep.For example, Bojarskaite et al report that global deletion of astrocyte IP3R2 increased microarousals and reduced average duration of NREM sleep^154,155^. In another study, genetic ablation of STIM1, an important protein involved in replenishing endoplasmic reticulum Ca^2+^, had no effect on baseline sleep^155^.

This discrepancy may be explained by differences in the sex of the animal models and in the regional specificity of astrocyte manipulation. Specifically, Bojarskaite and Ingiosi employed transgenic models that decreased astrocyte activity across the entire brain, while we specifically target astrocytes in the MnPO^154,155^. It is likely that astrocytes in different brain regions differentially affect sleep, and global manipulation of astrocytes may result in summative interactions that produce results dissimilar from manipulations in specific brain regions. Further, the Bojarskaite study exclusively used male mice, while we use female rats^154^. Interestingly, our findings in male rats (discussed in the “Sex differences” section below) more closely match the findings of the Bojarskaite study.

#### Astrocytes and Homeostatic Sleep Pressure

Extended periods of wakefulness are typically associated with increased sleep pressure, implying that astrocyte mediated enhancements in wakefulness should be accompanied by an increase in sleep pressure. However, despite increasing wake, Gq activation in MnPO astrocyte decreased cortical markers of homeostatic sleep pressure including NREM-SWA. To clarify the role of MnPO astrocytes in homeostatic sleep need, we sleep deprived animals for 6 hours, beginning at the end of the dark phase when sleep need is already high. We found that SD increased sleep in both control and p130PH animals during the first 6 hours of the light phase following SD. However, in the subsequent dark phase, SD did not increase sleep in p130PH animals. This was likely due to a ceiling effect, as just expressing p130PH in MnPO astrocytes increased dark phase sleep pressure to the same extent as 6 hours of sleep deprivation.

### Sex differences in MnPO astrocyte effects on Sleep

Our preliminary findings show that astrocytes in the MnPO produce different effects on sleep in male and female rats. P130PH expression in MnPO astrocytes increased sleep in female rats but showed a trend towards decreasing sleep in male rats.

These disparities may stem from (1) sex differences in the morphology and structure of astrocytes in the hypothalamus which may precipitate functional differences, sex differences in the number/type of neurons or gliotransmitter receptors in the MnPO. Early exposure to gonadal steroids permanently changes astrocyte architecture in the preoptic area^163^. Specifically, male astrocytes are more numerous and display greater branching of astrocytic processes than female astrocytes. As a result, male astrocytes may contact more synapses. These structural differences may be accompanied by distinct genetic profiles and may give rise to disparate effects on neurotransmission and downstream behaviors. For example, the AST2 subclass of astrocytes is enriched in genes involved in GABAergic transmission, while AST3 astrocytes primarily express genes for glutamatergic transmission^149^.

Because we OVXed female rats, the observed sex differences in astrocyte effects on sleep are likely a result of E2 mediated pre-programming of astrocytes during the prenatal stage, and not a product of current circulating hormones.

### E2 and Sleep

P130PH expression in MnPO astrocytes did not block E2’s sleep suppressing effects. Nevertheless, this does not discount a role for MnPO astrocytes in mediating E2’s effects. P130PH specifically reduces IP3 mediated Ca2+ release by preventing IP3 from binding to its receptor on the endoplasmic reticulum (ER)^131^. However, E2 may directly interact with estrogen receptors on the ER to release Ca^2+,^ bypassing the IP3 cascade. Thus, IP3 independent Ca^2+^ signaling in astrocytes may still mediate E2’s wake promoting effects

Finally, E2 may suppress oligodendrocyte release of prostaglandin D2 (PGD2), a potent sleep-promoting neuro-modulator, in the MnPO. Supporting this hypothesis, E2 has been shown to suppress PGD2 release in the VLPO, a NREM promoting center in the brain, which may lead to increased arousal^158^.

### We propose the following model that integrates our findings with existing knowledge of sleep-wake neurocircuitry

MnPO astrocytic Gq activation may be stimulated by inputs from wake-promoting nuclei, leading to tonic release of gliotransmitters that suppress the activity of sleep-promoting MnPO neurons. Supporting this hypothesis, the MnPO receives projections from wake-promoting structures such as the noradrenergic locus coeruleus and cholinergic basal forebrain. Further, norepinephrine and acetylcholine are

Our data suggests that MnPO astrocytes typically known to powerfully activates the astrocyte Gq-IP_3_-Ca^2+^ cascade, and astrocyte activity peaks in the dark phase when circulating levels of these wake-related neurotransmitters are high^150–152^. Hence, astrocytes activated by wake-promoting inputs may tonically release a gliotransmitter that inhibits sleep-promoting neurons. This gliotransmitter may be PGE2, adenosine or glutamate, each of which has been shown to suppress neuronal activity and/or enhance wake when introduced to the preoptic area.

### Clinical Significance of Astrocyte effects on Sleep

Women are disproportionately affected by insomnia. There are few reliable therapeutics for the treatment of insomnia. Benzodiazepine hypnotics and the “z” drugs (i.e. zolpidem, zaleplon,) are current leading therapeutics for insomnia. Unfortunately, while these medications promote sleep, they also suppress NREM delta power, a marker of deep restorative sleep that also plays a crucial role in memory consolidation^164,165^. Thus, “z” drugs promote light sleep that may not be restorative. In addition, these drugs exhibit negative effects on cognitive indices and may increase risk for dementia, which aligns with their inhibitory effect on NREM delta power^166^. Our work suggests that attenuation of astrocytes in the MnPO enhances sleep ***and*** NREM delta power in female rats. ***<u>These findings highlight astrocytes as a new therapeutic target for the treatment of insomnia and may contribute to the discovery of treatment options that enhance deep restorative sleep</u>***. However, any such therapeutic would need to be targeted to the MnPO using nanoparticles, as astrocytes in other brain regions may have a different effect on homeostatic sleep.

Sleep disturbances and derangements in astrocyte Ca^2+^ signaling are associated with neurodegenerative conditions including Huntington’s, Parkinsons, Alzheimer’s, Multiple Sclerosis and ALS^4,66–68,145,167–170^. In several of these conditions, sleep disturbances appear early in the disease process. For example, in Alzheimer’s disease, sleep disturbances appear decades before symptoms of cognitive decline are evident. It is not clear whether these sleep disturbances are prodromal, or causative for Alzheimer’s. Interestingly, GLP-1 agonists (i.e. Ozempic, liraglutide) have recently been shown to slow cognitive symptoms in neurodegenerative conditions^171–176^. Strikingly, this effect is mediated by GLP-1 agonist effects on astrocytes. However, no studies exist evaluating the effect of the GLP-1 drug class on sleep. Considering the emerging role of astrocytes in sleep, and the established role of astrocytes and sleep in neurodegeneration, it may be worth investigating whether these medications ameliorate sleep disturbances.

Aberrations in astrocyte Ca^2+^ signaling are also evident in chronic pain. Interestingly, there is a reciprocal connection between pain and sleep. Specifically, pain often leads to sleep disturbances, and sleep deprivation enhances sensitivity to pain^177,178^. The MnPO sends, and receives projections from the PB, an important nucleus implicated in pain and arousal. Recent findings suggest that PB neurons mediate pain-related awakenings from sleep^177^. Considering the significant role of astrocytes in chronic pain, it may be worth exploring whether PB projections to the MnPO are specifically implicated in pain-related arousals, and whether MnPO astrocytes are involved in this process. Interestingly, A2A antagonist infusion into the MnPO blocks the enhancing effect of SD on postoperative pain, potentially highlighting a role for astrocyte-derived adenosine in pain related awakenings.

## Materials and Methods

### Animals

Adult Long Evans female (200-350g) and male (250-350g) rats were obtained from Charles River Laboratories. Animals were housed in the Laboratory Animal Facilities at the University of Maryland, Baltimore, and maintained on a 12/12 dark:light cycle with *ad-libitum* access to food and water. Treatment times were based on zeitgeber time (ZT), with ZT 0 set to lights off. Due to resource constraints (number of telemetry transmitters and recording space), all experiments were run in multiple cohorts with a minimum of 2 animals per experimental group in each cohort. All experimental procedures were performed in accordance with the guidelines of the University of Maryland Institutional Animal Care and Use Committee. All procedures were in accordance with the National Institute of Health guide for the care and use of laboratory animals.

### Surgeries (4 pt space before)

Animal preparation and perioperative care/support were conducted in accordance with UMB IACUC protocols for all surgical procedures. All surgeries were aseptic and performed under isoflurane anesthesia with sterile materials.

For experiments where EEG/EMG data were collected, animals underwent the following: (1) a craniotomy to create burr holes for telemetry transmitter lead placement and viral infusion (2) stereotaxic AAV viral injection Ovariectomy (females) or Sham castration (males) and (4) intraperitoneal placement of a bi-potential telemetry transmitter (DSI, St. Paul, Minn). For animals in the p130PH study, all surgical procedures (#1-#4) took place in one single survival surgery. Briefly, a midline incision was made along the dorsal head and neck. 3 burr holes were drilled in the skull: one at Bregma for stereotaxic viral infusion into the MnPO, and 2 for telemetry lead placement (AP +2mm and ML +1.5mm; AP -7mm and ML -1.5mm). AAV virus was injected into the MnPO at the appropriate stereotaxic coordinates (Bregma, dorsoventral -7.2mm) using a Hamilton syringe. After viral infusion, a midline skin incision was made along the back followed by incisions along the flank abdominal muscles. Abdominal wall incisions were bilateral for females, and unilateral for males. For females, the ovarian fat pads were visualized using the bilateral incisions and both ovaries were removed. For males and females, telemetry transmitters were placed in the intraperitoneal space through the abdominal wall incisions and secured to the internal abdominal wall. The transmitter leads were subcutaneously threaded along the back to the head incision. EEG leads were wrapped around stainless steel screws implanted in the remaining 2 burr holes in the skull and secured in place using dental acrylic. For males, Sham castration took place before the stereotaxic components of the surgery.

For animals in the DREADD study, the viral infusion took place in a separate surgery at least 15 days before the OVX/Sham castration + telemetry transmitter implant surgery. All animals received post-surgical monitoring and analgesia according to IACUC protocols. All animals were allowed to recover for at least 10 days after their final surgery before experiments.

Animals in the P130 study underwent a single surgery involving viral infusion, OVX, and telemeter implant.

For immunohistochemical studies, female rodents were used and underwent the following surgical procedure in one survival surgery: (1) OVX. OVX took place as described above.

### Data Acquisition (4 pt space before)

EEG and EMG data were continuously collected by telemetry transmitters (DSI, St. Paul, Minn) implanted in each animal and relayed to Ponemah software (DSI, St. Paul, Minn) via receiver bases placed under each animal’s home cage. Vigilance states were scored in 10-second epochs by visual inspection of EEG/EMG signals using Neuroscore (DSI, St. Paul, Minn). Epochs were scored as wake, NREM or REM according to the paradigm described in Table 1 below. The scored data were used to calculate total time in state, number of bouts, and average bout duration for wake, NREM and REM.

### Power Spectral Density (PSD) and NREM-SWA (4 pt space before)

The power spectral density (PSD) is a visual representation of how much power an EEG signal contains across a range of frequencies. This can provide information about various frequencies of interest, including theta (4-8Hz) and SWA (0.5-4Hz) which have been implicated in memory consolidation and sleep propensity/sleep need respectively. For each vigilance state, a fast Fourier transform was used to compute the PSD from 0.5-10Hz in 0.25Hz stepwise bins. Each bin was normalized to the mean total power (averaged across wake, NREM and REM) for the entire 24-hour period of the same day or a baseline day.

NREM-SWA activity (a measure of homeostatic sleep drive) was calculated by computing the EEG power specifically in the delta band (0.5-4Hz) of NREM bouts at hourly intervals from ZT0 to ZT23. Delta band power was normalized to mean total power averaged across the three vigilance states for the full 24-hour period.

For Gi experiments, PSD were generated for the full 12hr phase during which Saline/CNO injections took place, and for the first 4 hours of that phase. For p130PH experiment, PSD were generated for each phase (Light and Dark Phase).

NREM-SWA was assessed by quantifying EEG spectral power in the delta band (0.5-4Hz) in hourly bins and normalizing to the 24h total power for each experimental day. For Gi experiments, NREM-SWA was visualized for the first 6 hours of the phase to determine acute treatment effects, and the last 6 hours of the phase to evaluate for rebound effects. For p130PH experiments, NREM-SWA was presented for the full light or dark phase.

### Immunohistochemistry (4 pt space before)

Animals were deeply anesthetized with isoflurane and transcardially perfused with a 0.9% saline + 2% sodium nitrite solution followed by 4% paraformaldehyde in 0.05M KPBS. Brains were extracted and immediately post-fixed in 4% paraformaldehyde for 24 hours. Subsequently, brains were cryoprotected in 30% sucrose solution, flash frozen on dry ice and stored at -80C. A cryostat was used to section each brain into 4 replicate series with 30 micron thick slices cut along the coronal plane. The brain slices were stored free floating in an ethylene-glycol and PVP based solution at -20°C while awaiting processing.

For immunohistochemical analysis, the sections were rinsed with PBS and incubated in blocking solution for 1 hour. Sections were then incubated with primary antibodies for the protein of interest for 24 hours at 4C. Next, sections were washed in PBS and incubated with a fluorescently labelled secondary antibody for 2 hours at room temperature. Following this, the sections were rinsed in PBS to remove excess secondary antibody, incubated with DAPI for 3 minutes to visualize cell nuclei, and rinsed with PBS a final time. Finally, slices were mounted on Superfrost slides and coverslipped.

### Experimental Protocols (4 pt space before)

#### Using Gi-DREADD to decrease astrocyte activity

To determine the effects of decreasing astrocyte activity on sleep and cortical markers of homeostatic sleep propensity, we used AAV5-GFAP-hM4DGi-mCherry to selectively express Gi-DREADD in MnPO astrocytes of adult female rats (n=5). Animals were ovariectomized, intraperitoneally implanted with telemetry transmitters, and randomly assigned to one of two treatment groups in a within-animal double cross-over design.

To determine astrocyte effects on homeostatic sleep in the Dark Phase, animals were treated with Saline Vehicle or clozapine-N-oxide, a ligand for Gi-DREADD at the start of the Dark Phase (ZT0). Animals were also treated with Sesame Oil Control at the time of these injections, followed by 2 consecutive injections of E2 separated by 24 hours. In this dissertation, we only present the results of the Saline/CNO treatments on the Oil Day, with Gi-DREADD/E2 data presented in the Appendix. After a wash out period of at least 5 days, animals received the opposite treatment at the start of the Dark Phase (ZT0); animals that initially received Saline Vehicle now received CNO and vice versa.

Following another washout period of at least 5 days, we next examined astrocyte effects on sleep in the Light Phase. Animals were randomly assigned to two groups, receiving either Saline or CNO at the start of the Light Phase (ZT12). After a minimum washout period of 5 days, animals entered the opposite treatment arm.

EEG/EMG signals were collected for the duration of these experiments except during the wash out periods. Finally, animals remained in their home cages for the duration of this experiment.

#### Using p130PH to decrease astrocyte activity

To evaluate the effects of decreasing astrocyte activity on sleep, we constitutively expressed p130PH in MnPO astrocytes of adult female rats. To do this, we injected AAV5-gfaaBC1D-p130PH-mRFP into the MnPO or AAV5-gfaABC1D-tdTomato as a control. Animals were OVX and implanted with a telemetry transmitter during the same surgery. After a minimum 10 day recovery period, animal sleep was recorded for at-least 24 hours. Animals remained in their home cage for the duration of this experiment.

#### Evaluating whether p130PH expression in astrocytes will block E2’s effects on Sleep

To examine whether decreasing astrocyte activity blocks E2’s effects on sleep, we constitutively expressed p130PH in MnPO astrocytes of adult female rats. To do this, we injected AAV5-gfaaBC1D-p130PH-mRFP into the MnPO or AAV5-gfaABC1D-tdTomato as a control. Animals were ovariectomized and intraperitoneally implanted with telemetry transmitters to collect sleep data.

Animals were treated with Sesame Oil Control at the start of the Dark Phase (ZT0), followed by a subcutaneous E2 injection (5uq) 24 hours later, and a final E2 injection (10ug) 24 hours after that. EEG/EMG signals were collected until 48 hours after the final E2 injection. The Oil day and Post E2 day were scored and evaluated. Animals remained in their home cages for the duration of this experiment.

#### Evaluating whether p130PH expression in astrocytes has different effects on male sleep

To determine the effects of decreasing astrocyte activity on sleep, we constitutively expressed p130PH in MnPO astrocytes of adult male rats. To do this, we injected AAV5-gfaaBC1D-p130PH-mRFP into the MnPO or AAV5-gfaABC1D-tdTomato as a control. Animals were Sham castrated and implanted with a telemetry transmitter during the same surgery. After a minimum 10 day recovery period, animal sleep was recorded for at-least 24 hours. Animals remained in their home cage for the duration of this experiment

#### Sleep Deprivation

To clarify the contribution of astrocytes to markers of homeostatic sleep drive, we sleep deprived animals for 6 hours starting at the last hour of the dark phase (ZT11) or the first hour of the light phase (ZT12). We used adult female rats expressing p130PH in MnPO astrocytes, or tDTomato as a control. Animals were also OVX and implanted with telemetry transmitters to collect sleep data.

Animals were randomly assigned to two groups: undisturbed sleep or sleep deprived. SD began at ZT11 or ZT12, and gentle handling was used to keep animals awake. After 6 hours of SD, animals were allowed undisturbed recovery sleep for 18 hours. Animals in the US group were allowed undisturbed sleep for the full 24 hour experimental day. After a minimum 5 day recovery period, animals crossed over into the opposite treatment arm. Animals remained in their home cages for the duration of the experiment.

### Statistics

For Gi-DREADD experiments, 4 hour totals for time in state, number of bouts and average bout duration are presented as mean +/-SD and analyzed using paired t-test. For p130PH experiments, 12 hour totals for time in state, number of bouts and average bout durations are presented as mean +/-SEM and analyzed using unpaired t-test. Hourly vigilance state time-course plots, PSD and NREM-SWA are presented as mean +/-SEM and analyzed with Mixed Effects Analysis or Repeated Measures 2 -way ANOVA. Statistical tests were conducted using GraphPad Prism.

For p130PH experiments, 12 hour totals for time in state, number of bouts and average bout durations are presented as mean +/-SEM and analyzed using unpaired t-test. Hourly vigilance state time-course plots, PSD and NREM-SWA are presented as mean +/-SEM and analyzed with Mixed Effects Analysis or Repeated Measures 2-way ANOVA. Statistical tests were conducted using GraphPad Prism.

## Notes

### Competing Interest Statement

The authors have declared no competing interest.

### Summary of Updates

The Funding Source has been corrected to reflect the accurate grant number.

### Funding

- NIH/NHLBI: R01HL85037

## References

1. Xie L, Kang H, Xu Q, et al. Sleep drives metabolite clearance from the adult brain. Science (1979). 2013;342(6156). doi:10.1126/science.1241224

2. Okun ML, Mancuso RA, Hobel CJ, Schetter CD, Coussons-Read M. Poor sleep quality increases symptoms of depression and anxiety in postpartum women. J Behav Med. 2018;41(5). doi:10.1007/s10865-0189950-7

3. Kim KM, Yang KI. Sleep and Epilepsy. Sleep Med Res. 2023;14(2). doi:10.17241/smr.2023.01781

4. Olsson M, Ärlig J, Hedner J, Blennow K, Zetterberg H. Sleep deprivation and cerebrospinal fluid biomarkers for Alzheimer’s disease. Sleep. 2018;41(5). doi:10.1093/sleep/zsy025

5. Shan Z, Ma H, Xie M, et al. Sleep duration and risk of type 2 diabetes: A meta-analysis of prospective studies. Diabetes Care. 2015;38(3). doi:10.2337/dc14-2073

6. Harding K, Feldman M. Sleep Disorders and Sleep Deprivation: An Unmet Public Health Problem. J Am Acad Child Adolesc Psychiatry. 2008;47(4). doi:10.1097/01.chi.0000270812.55636.3b

7. Liu Y, Wheaton AG, Chapman DP, Cunningham TJ, Lu H, Croft JB. Prevalence of Healthy Sleep Duration among Adults — United States, 2014. MMWR Morb Mortal Wkly Rep. 2016;65(6). doi:10.15585/mmwr.mm6506a1

8. Phillips AJK, Robinson PA, Kedziora DJ, Abeysuriya RG. Mammalian sleep dynamics: How diverse features arise from a common physiological framework. PLoS Comput Biol. 2010;6(6). doi:10.1371/journal.pcbi.1000826

9. Miyazaki S, Liu CY, Hayashi Y. Sleep in vertebrate and invertebrate animals, and insights into the function and evolution of sleep. Neurosci Res. 2017;118. doi:10.1016/j.neures.2017.04.017

10. Dement W. The occurence of low voltage, fast, electroencephalogram patterns during behavioral sleep in the cat. Electroencephalogr Clin Neurophysiol. 1958;10(2). doi:10.1016/0013-4694(58)90037-3

11. Campbell SS, Tobler I. Animal sleep: A review of sleep duration across phylogeny. Neurosci Biobehav Rev. 1984;8(3). doi:10.1016/0149-7634(84)90054-X

12. Dijk DJ. Regulation and functional correlates of slow wave sleep. Journal of Clinical Sleep Medicine. 2009;5(2 SUPPL.). doi:10.5664/jcsm.5.2s.s6

13. Berry RB, Albertario CL, Harding SM, et al. The AASM Manual for the Scoring of Sleep and Associated Events. Journal of Clinical Sleep Medicine. 2018;2.5.

14. Feinberg I, Floyd TC. Systematic Trends Across the Night in Human Sleep Cycles. Psychophysiology. 1979;16(3). doi:10.1111/j.1469-8986.1979.tb02991.x

15. Mendoza J. Nighttime Light Hurts Mammalian Physiology: What Diurnal Rodent Models Are Telling Us. Clocks Sleep. 2021;3(2). doi:10.3390/clockssleep3020014

16. Zhang B, Wing YK. Sex differences in insomnia: A meta-analysis. Sleep. 2006;29(1). doi:10.1093/sleep/29.1.85

17. Suh S, Cho N, Zhang J. Sex Differences in Insomnia: from Epidemiology and Etiology to Intervention. Curr Psychiatry Rep. 2018;20(9). doi:10.1007/s11920-018-0940-9

18. Shaib F, Attarian H. Sex and gender differences in sleep disorders: An overview. In: Principles of Gender-Specific Medicine: Sex and Gender-Specific Biology in the Postgenomic Era. ; 2023. doi:10.1016/B978-0-323-88534-8.00036-5

19. Makarem N, Aggarwal B. Gender Differences in Associations between Insufficient Sleep and Cardiovascular Disease Risk Factors and Endpoints: A Contemporary Review. Gend Genome. 2017;1(2). doi:10.1089/gg.2017.0001

20. Cappuccio FP, Stranges S, Kandala NB, et al. Gender-specific associations of short sleep duration with prevalent and incident hypertension: The whitehall II study. Hypertension. 2007;50(4). doi:10.1161/HYPERTENSIONAHA.107.095471

21. Bixler EO, Papaliaga MN, Vgontzas AN, et al. Women sleep objectively better than men and the sleep of young women is more resilient to external stressors: Effects of age and menopause. J Sleep Res. 2009;18(2). doi:10.1111/j.1365-2869.2008.00713.x

22. Goel N, Kim H, Lao RP. Gender differences in polysomnographic sleep in young healthy sleepers. Chronobiol Int. 2005;22(5). doi:10.1080/07420520500263235

23. Roehrs T, Kapke A, Roth T, Breslau N. Sex differences in the polysomnographic sleep of young adults: A community-based study. Sleep Med. 2006;7(1). doi:10.1016/j.sleep.2005.05.008

24. Baker FC, Sassoon SA, Kahan T, et al. Perceived poor sleep quality in the absence of polysomnographic sleep disturbance in women with severe premenstrual syndrome. J Sleep Res. 2012;21(5). doi:10.1111/j.1365-2869.2012.01007.x

25. Vitiello M V., Larsen LH, Moe KE. Age-related sleep change: Gender and estrogen effects on the subjective-objective sleep quality relationships of healthy, noncomplaining older men and women. J Psychosom Res. 2004;56(5). doi:10.1016/S0022-3999(04)00023-6

26. Cui J, Shen Y, Li R. Estrogen synthesis and signaling pathways during aging: From periphery to brain. Trends Mol Med. 2013;19(3). doi:10.1016/j.molmed.2012.12.007

27. Milner TA, McEwen BS, Hayashi S, Li CJ, Reagan LP, Alves SE. Ultra-structural evidence that hippocampal alpha estrogen receptors are located at extranuclear sites. Journal of Comparative Neurology. 2001;429(3). doi:10.1002/1096-9861(20010115)429:3<355::aid-cne1>3.0.co;2-%23

28. Milner TA, Ayoola K, Drake CT, et al. Ultrastructural localization of estrogen receptor β immunoreactivity in the rat hippocampal formation. Journal of Comparative Neurology. 2005;491(2). doi:10.1002/cne.20724

29. Laredo SA, Villalon Landeros R, Trainor BC. Rapid effects of estrogens on behavior: Environmental modulation and molecular mechanisms. Front Neuroendocrinol. 2014;35(4). doi:10.1016/j.yfrne.2014.03.005

30. Qiu J, Rønnekleiv OK, Kelly MJ. Modulation of hypothalamic neuronal activity through a novel G-protein-coupled estrogen membrane receptor. Steroids. 2008;73(9-10). doi:10.1016/j.steroids.2007.11.008

31. Revankar CM, Cimino DF, Sklar LA, Arterburn JB, Prossnitz ER. A transmembrane intracellular estrogen receptor mediates rapid cell signaling. Science (1979). 2005;307(5715). doi:10.1126/science.1106943

32. Baker FC, Driver HS. Circadian rhythms, sleep, and the menstrual cycle. Sleep Med. 2007;8(6). doi:10.1016/j.sleep.2006.09.011

33. Sedov ID, Cameron EE, Madigan S, Tomfohr-Madsen LM. Sleep quality during pregnancy: A meta-analysis. Sleep Med Rev. 2018;38. doi:10.1016/j.smrv.2017.06.005

34. Parry BL, Mostofi N, Leveau B, et al. Sleep EEG studies during early and late partial sleep deprivation in premenstrual dysphoric disorder and normal control subjects. Psychiatry Res. 1999;85(2). doi:10.1016/S0165-1781(98)00128-0

35. De Zambotti M, Willoughby AR, Sassoon SA, Colrain IM, Baker FC. Menstrual cycle-related variation in physiological sleep in women in the early menopausal transition. Journal of Clinical Endocrinology and Metabolism. 2015;100(8). doi:10.1210/jc.2015-1844

36. Aukia L, Paavonen EJ, Jänkälä T, et al. Insomnia symptoms increase during pregnancy, but no increase in sleepiness - Associations with symptoms of depression and anxiety. Sleep Med. 2020;72. doi:10.1016/j.sleep.2020.03.031

37. Baker FC, Lampio L, Saaresranta T, Polo-Kantola P. Sleep and Sleep Disorders in the Menopausal Transition. Sleep Med Clin. 2018;13(3). doi:10.1016/j.jsmc.2018.04.011

38. Nelson HD. Menopause. The Lancet. 2008;371(9614). doi:10.1016/S0140-6736(08)60346-3

39. Baker FC, Mitchell D, Driver HS. Oral contraceptives alter sleep and raise body temperature in young women. Pflugers Arch. 2001;442(5). doi:10.1007/s004240100582

40. Baker FC, Waner JI, Vieira EF, Taylor SR, Driver HS, Mitchell D. Sleep and 24 hour body temperatures: A comparison in young men, naturally cycling women and women taking hormonal contraceptives. Journal of Physiology. 2001;530(3). doi:10.1111/j.1469-7793.2001.0565k.x

41. Hadjimarkou MM, Benham R, Schwarz JM, Holder MK, Mong JA. Estradiol suppresses rapid eye movement sleep and activation of sleep-active neurons in the ventrolateral preoptic area. European Journal of Neuroscience. 2008;27(7). doi:10.1111/j.1460-9568.2008.06142.x

42. Smith PC, Phillips DJ, Pocivavsek A, et al. Estradiol influences adenosinergic signaling and nonrapid eye movement sleep need in adult female rats. Sleep. 2022;45(3). doi:10.1093/sleep/zsab225

43. Triarhou LC. The percipient observations of Constantin von Economo on encephalitis lethargica and sleep disruption and their lasting impact on contemporary sleep research. Brain Res Bull. 2006;69(3). doi:10.1016/j.brainresbull.2006.02.002

44. Saper CB, Scammell TE, Lu J. Hypothalamic regulation of sleep and circadian rhythms. Nature. 2005;437(7063). doi:10.1038/nature04284

45. Saper CB, Chou TC, Scammell TE. The sleep switch: Hypothalamic control of sleep and wakefulness. Trends Neurosci. 2001;24(12). doi:10.1016/S0166-2236(00)02002-6

46. Arrigoni E, Fuller PM. The Sleep-Promoting Ventrolateral Preoptic Nucleus: What Have We Learned over the Past 25 Years? Int J Mol Sci. 2022;23(6). doi:10.3390/ijms23062905

47. Rothhaas R, Chung S. Role of the Preoptic Area in Sleep and Thermoregulation. Front Neurosci. 2021;15. doi:10.3389/fnins.2021.664781

48. Vanini G, Bassana M, Mast M, et al. Activation of Preoptic GABAergic or Glutamatergic Neurons Modulates Sleep-Wake Architecture, but Not Anesthetic State Transitions. Current Biology. 2020;30(5). doi:10.1016/j.cub.2019.12.063

49. Machado NLS, Todd WD, Kaur S, Saper CB. Median preoptic GABA and glutamate neurons exert differential control over sleep behavior. Current Biology. 2022;32(9). doi:10.1016/j.cub.2022.03.039

50. Alam MA, Kumar S, McGinty D, Alam MN, Szymusiak R. Neuronal activity in the preoptic hypothalamus during sleep deprivation and recovery sleep. J Neurophysiol. 2014;111(2). doi:10.1152/jn.00504.2013

51. Borbely AA, Tobler I. Endogenous sleep-promoting substances and sleep regulation. Physiol Rev. 1989;69(2). doi:10.1152/physrev.1989.69.2.605

52. Deboer T, Vansteensel MJ, Détári L, Meijer JH. Sleep states alter activity of suprachiasmatic nucleus neurons. Nat Neurosci. 2003;6(10). doi:10.1038/nn1122

53. Sanchez REA, Kalume F, de la Iglesia HO. Sleep timing and the circadian clock in mammals: Past, present and the road ahead. Semin Cell Dev Biol. 2022;126. doi:10.1016/j.semcdb.2021.05.034

54. Fisher SP, Foster RG, Peirson SN. The circadian control of sleep. Handb Exp Pharmacol. 2013;217. doi:10.1007/978-3-642-25950-0_7

55. Peng W, Wu Z, Song K, Zhang S, Li Y, Xu M. Regulation of sleep homeostasis mediator adenosine by basal forebrain glutamatergic neurons. Science (1979). 2020;369(6508). doi:10.1126/SCIENCE.ABB0556

56. Blanco-Centurion C, Xu M, Murillo-Rodriguez E, et al. Adenosine and sleep homeostasis in the basal forebrain. Journal of Neuroscience. 2006;26(31). doi:10.1523/JNEUROSCI.2181-06.2006

57. Nakamura TJ, Moriya T, Inoue S, et al. Estrogen differentially regulates expression of Per1 and Per2 genes between central and peripheral clocks and between reproductive and nonreproductive tissues in female rats. J Neurosci Res. 2005;82(5). doi:10.1002/jnr.20677

58. Smith PC, Cusmano DM, Viechweg SS, Mong JA. Estradiol Action at the Median Preoptic Nucleus is Necessary and Sufficient for 1 Sleep Suppression in Female Rats 2. doi:10.1101/2020.07.29.223669

59. Methippara MM, Kumar S, Alam N, Szymusiak R, McGinty D. Effects on sleep of microdialysis of adenosine A1 and A 2a receptor analogs into the lateral preoptic area of rats. Am J Physiol Regul Integr Comp Physiol. 2005;289(6 58-6). doi:10.1152/ajpregu.00247.2005

60. Hines DJ, Haydon PG. Astrocytic adenosine: From synapses to psychiatric disorders. Philosophical Transactions of the Royal Society B: Biological Sciences. 2014;369(1654). doi:10.1098/rstb.2013.0594

61. Bjorness TE, Dale N, Mettlach G, et al. An adenosine-mediated glial-neuronal circuit for homeostatic sleep. Journal of Neuroscience. 2016;36(13). doi:10.1523/JNEUROSCI.3906-15.2016

62. Acaz-Fonseca E, Sanchez-Gonzalez R, Azcoitia I, Arevalo MA, Garcia-Segura LM. Role of astrocytes in the neuroprotective actions of 17β-estradiol and selective estrogen receptor modulators. Mol Cell Endocrinol. 2014;389(1-2). doi:10.1016/j.mce.2014.01.009

63. Benarroch EE. Neuron-astrocyte interactions: Partnership for normal function and disease in the central nervous system. In: Mayo Clinic Proceedings. Vol 80. ; 2005. doi:10.4065/80.10.1326

64. Oberheim NA, Goldman SA, Nedergaard M. Heterogeneity of astrocytic form and function. Methods in Molecular Biology. 2012;814. doi:10.1007/978-1-61779-452-0_3

65. Vasile F, Dossi E, Rouach N. Human astrocytes: structure and functions in the healthy brain. Brain Struct Funct. 2017;222(5). doi:10.1007/s00429-017-1383-5

66. Yun SP, Kam TI, Panicker N, et al. Block of A1 astrocyte conversion by microglia is neuroprotective in models of Parkinson’s disease. Nat Med. 2018;24(7). doi:10.1038/s41591-018-0051-5

67. Tong X, Ao Y, Faas GC, et al. Astrocyte Kir4.1 ion channel deficits contribute to neuronal dysfunction in Huntington’s disease model mice. Nat Neurosci. 2014;17(5). doi:10.1038/nn.3691

68. Khakh BS, Beaumont V, Cachope R, Munoz-Sanjuan I, Goldman SA, Grantyn R. Unravelling and Exploiting Astrocyte Dysfunction in Huntington’s Disease. Trends Neurosci. 2017;40(7). doi:10.1016/j.tins.2017.05.002

69. Siracusa R, Fusco R, Cuzzocrea S. Astrocytes: Role and functions in brain pathologies. Front Pharmacol. 2019;10(SEP). doi:10.3389/fphar.2019.01114

70. Weis WI, Kobilka BK. The Molecular Basis of G Protein-Coupled Receptor Activation. Annu Rev Biochem. 2018;87. doi:10.1146/annurevbiochem-060614-033910

71. Cabrera-Vera TM, Vanhauwe J, Thomas TO, et al. Insights into G Protein Structure, Function, and Regulation. Endocr Rev. 2003;24(6). doi:10.1210/er.2000-0026

72. Durkee CA, Covelo A, Lines J, Kofuji P, Aguilar J, Araque A. G i/o protein-coupled receptors inhibit neurons but activate astrocytes and stimulate gliotransmission. Glia. 2019;67(6). doi:10.1002/glia.23589

73. D’Ascenzo M, Fellin T, Terunuma M, et al. mGluR5 stimulates gliotransmission in the nucleus accumbens. Proc Natl Acad Sci U S A. 2007;104(6). doi:10.1073/pnas.0609408104

74. Abbracchio MP, Ceruti S. Roles of P2 receptors in glial cells: Focus on astrocytes. In: Purinergic Signalling. Vol 2. ; 2006. doi:10.1007/s11302006-9016-0

75. Gordon GRJ, Baimoukhametova D V., Hewitt SA, Rajapaksha WRAKJS, Fisher TE, Bains JS. Norepinephrine triggers release of glial ATP to increase postsynaptic efficacy. Nat Neurosci. 2005;8(8). doi:10.1038/nn1498

76. Navarrete M, Díez A, Araque A. Astrocytes in endocannabinoid signalling. Philosophical Transactions of the Royal Society B: Biological Sciences. 2014;369(1654). doi:10.1098/rstb.2013.0599

77. Boison D, Chen JF, Fredholm BB. Adenosine signaling and function in glial cells. Cell Death Differ. 2010;17(7). doi:10.1038/cdd.2009.131

78. Hertz L, Lovatt D, Goldman SA, Nedergaard M. Adrenoceptors in brain: Cellular gene expression and effects on astrocytic metabolism and [Ca2+]i. Neurochem Int. 2010;57(4). doi:10.1016/j.neuint.2010.03.019

79. Morin D, Sapena R, Zini R, Onteniente B, Tillement JP. Characterization of β-adrenergic receptors of freshly isolated astrocytes and neurons from rat brain. Life Sci. 1996;60(4-5). doi:10.1016/S0024-3205(96)00632-7

80. Xin W, Schuebel KE, Jair K wing, et al. Ventral midbrain astrocytes display unique physiological features and sensitivity to dopamine D2 receptor signaling. Neuropsychopharmacology. 2019;44(2). doi:10.1038/s41386-018-0151-4

81. Nam MH, Han KS, Lee J, et al. Activation of Astrocytic µ-Opioid Receptor Causes Conditioned Place Preference. Cell Rep. 2019;28(5). doi:10.1016/j.celrep.2019.06.071

82. Mariotti L, Losi G, Sessolo M, Marcon I, Carmignoto G. The inhibitory neurotransmitter GABA evokes long-lasting Ca2+ oscillations in cortical astrocytes. Glia. 2016;64(3). doi:10.1002/glia.22933

83. Jennings A, Tyurikova O, Bard L, et al. Dopamine elevates and lowers astroglial Ca2+ through distinct pathways depending on local synaptic circuitry. Glia. 2017;65(3). doi:10.1002/glia.23103

84. Gould T, Chen L, Emri Z, et al. GABAB receptor-mediated activation of astrocytes by gamma-hydroxybutyric acid. Philosophical Transactions of the Royal Society B: Biological Sciences. 2014;369(1654). doi:10.1098/rstb.2013.0607

85. Copeland CS, Wall TM, Sims RE, et al. Astrocytes modulate thalamic sensory processing via mGlu2 receptor activation. Neuropharmacology. 2017;121. doi:10.1016/j.neuropharm.2017.04.019

86. Hua X, Malarkey EB, Sunjara V, Rosenwald SE, Li WH, Parpura V. Ca2+-Dependent Glutamate Release Involves Two Classes of Endoplasmic Reticulum Ca2+ Stores in Astrocytes. J Neurosci Res. 2004;76(1). doi:10.1002/jnr.20061

87. Araque A, Li N, Doyle RT, Haydon PG. SNARE protein-dependent glutamate release from astrocytes. Journal of Neuroscience. 2000;20(2). doi:10.1523/jneurosci.20-02-00666.2000

88. Murat CDB, García-Cáceres C. Astrocyte gliotransmission in the regulation of systemic metabolism. Metabolites. 2021;11(11). doi:10.3390/metabo11110732

89. Parpura V, Basarsky TA, Liu F, Jeftinija K, Jeftinija S, Haydon PG. Glutamate-mediated astrocyte-neuron signalling. Nature. 1994;369(6483). doi:10.1038/369744a0

90. Orellana JA, Froger N, Ezan P, et al. ATP and glutamate released via astroglial connexin 43 hemichannels mediate neuronal death through activation of pannexin 1 hemichannels. J Neurochem. 2011;118(5). doi:10.1111/j.1471-4159.2011.07210.x

91. Ye ZC, Wyeth MS, Baltan-Tekkok S, Ransom BR. Functional hemichannels in astrocytes: A novel mechanism of glutamate release. Journal of Neuroscience. 2003;23(9). doi:10.1523/jneurosci.23-09-03588.2003

92. Chever O, Lee CY, Rouach N. Astroglial connexin43 hemichannels tune basal excitatory synaptic transmission. Journal of Neuroscience. 2014;34(34). doi:10.1523/JNEUROSCI.0015-14.2014

93. Li D, Ropert N, Koulakoff A, Giaume C, Oheim M. Lysosomes are the major vesicular compartment undergoing Ca 2+-regulated exocytosis from cortical astrocytes. Journal of Neuroscience. 2008;28(30). doi:10.1523/JNEUROSCI.0744-08.2008

94. Zhang Z, Chen G, Zhou W, et al. Regulated ATP release from astrocytes through lysosome exocytosis. Nat Cell Biol. 2007;9(8). doi:10.1038/ncb1620

95. Woo DH, Han KS, Shim JW, et al. TREK-1 and Best1 Channels Mediate Fast and Slow Glutamate Release in Astrocytes upon GPCR Activation. Cell. 2012;151(1). doi:10.1016/j.cell.2012.09.005

96. Brancaccio M, Patton AP, Chesham JE, Maywood ES, Hastings MH. Astrocytes Control Circadian Timekeeping in the Suprachiasmatic Nucleus via Glutamatergic Signaling. Neuron. 2017;93(6). doi:10.1016/j.neuron.2017.02.030

97. Halassa MM, Florian C, Fellin T, et al. Astrocytic Modulation of Sleep Homeostasis and Cognitive Consequences of Sleep Loss. Neuron. 2009;61(2). doi:10.1016/j.neuron.2008.11.024

98. Tso MCF, Herzog ED. Was Cajal right about sleep? BMC Biol. 2015;13(1). doi:10.1186/s12915-015-0178-5

99. Foley J, Blutstein T, Lee S, Erneux C, Halassa MM, Haydon P. Astrocytic IP3/Ca2+ signaling modulates theta rhythm and REM sleep. Front Neural Circuits. 2017;11. doi:10.3389/fncir.2017.00003

100. Ingiosi AM, Hayworth CR, Frank MG. Activation of Basal Forebrain Astrocytes Induces Wakefulness without Compensatory Changes in Sleep Drive. Journal of Neuroscience. 2023;43(32). doi:10.1523/JNEUROSCI.0163-23.2023

101. Cai P, Huang SN, Lin ZH, et al. Regulation of wakefulness by astrocytes in the lateral hypothalamus. Neuropharmacology. 2022;221. doi:10.1016/j.neuropharm.2022.109275

102. Liu PC, Yao W, Chen XY, et al. Parabrachial nucleus astrocytes regulate wakefulness and isoflurane anesthesia in mice. Front Pharmacol. 2023;13. doi:10.3389/fphar.2022.991238

103. Tsunematsu T, Sakata S, Sanagi T, Tanaka KF, Matsui K. Region-Specific and State-Dependent Astrocyte Ca21 Dynamics during the SleepWake Cycle in Mice. Journal of Neuroscience. 2021;41(25). doi:10.1523/JNEUROSCI.2912-20.2021

104. Ingiosi AM, Frank MG. Noradrenergic Signaling in Astrocytes Influences Mammalian Sleep Homeostasis. Clocks Sleep. 2022;4(3). doi:10.3390/clockssleep4030028

105. Ingiosi AM, Hayworth CR, Harvey DO, et al. A Role for Astroglial Calcium in Mammalian Sleep and Sleep Regulation. Current Biology. 2020;30(22). doi:10.1016/j.cub.2020.08.052

106. Peng W, Liu X, Ma G, et al. Adenosine-independent regulation of the sleep–wake cycle by astrocyte activity. Cell Discov. 2023;9(1). doi:10.1038/s41421-022-00498-9

107. Kurogi Y, Sanagi T, Ono D, Tsunematsu T. Chemogenetic activation of astrocytes modulates sleep–wakefulness states in a brain region-dependent manner. SLEEP Advances. 2024;5(1):91. doi:10.1093/SLEEPADVANCES/ZPAE091

108. Kim JH, Choi IS, Jeong JY, Jang IS, Lee MG, Suk K. Astrocytes in the ventrolateral preoptic area promote sleep. Journal of Neuroscience. 2020;40(47). doi:10.1523/JNEUROSCI.1486-20.2020

109. Patton AP, Smyllie NJ, Chesham JE, Hastings MH. Astrocytes Sustain Circadian Oscillation and Bidirectionally Determine Circadian Period, But Do Not Regulate Circadian Phase in the Suprachiasmatic Nucleus. Journal of Neuroscience. 2022;42(28). doi:10.1523/JNEUROSCI.2337-21.2022

110. Vanderheyden WM, Goodman AG, Taylor RH, Frank MG, Van Dongen HPA, Gerstner JR. Astrocyte expression of the Drosophila TNF-alpha homologue, Eiger, regulates sleep in flies. PLoS Genet. 2018;14(10). doi:10.1371/journal.pgen.1007724

111. Haydon PG. Astrocytes and the modulation of sleep. Curr Opin Neurobiol. 2017;44. doi:10.1016/j.conb.2017.02.008

112. Collado P, Beyer C, Hutchison JB, Holman SD. Hypothalamic distribution of astrocytes is gender-related in Mongolian gerbils. Neurosci Lett. 1995;184(2). doi:10.1016/0304-3940(94)11175-I

113. Johnson RT, Breedlove SM, Jordan CL. Sex differences and laterality in astrocyte number and complexity in the adult rat medial amygdala. Journal of Comparative Neurology. 2008;511(5). doi:10.1002/cne.21859

114. Mong JA, Glaser E, McCarthy MM. Gonadal steroids promote glial differentiation and alter neuronal morphology in the developing hypothalamus in a regionally specific manner. Journal of Neuroscience. 1999;19(4). doi:10.1523/jneurosci.19-04-01464.1999

115. Weisz J, Ward IL. Plasma testosterone and progesterone titers of pregnant rats, their male and female fetuses, and neonatal offspring. Endocrinology. 1980;106(1). doi:10.1210/endo-106-1-306

116. Mong JA, Kurzweil RL, Davis AM, Rocca MS, McCarthy MM. Evidence for sexual differentiation of glia in rat brain. Horm Behav. 1996;30(4). doi:10.1006/hbeh.1996.0058

117. Mong JA, Blutstein T. Estradiol modulation of astrocytic form and function: Implications for hormonal control of synaptic communication. Neuroscience. 2006;138(3). doi:10.1016/j.neuroscience.2005.10.017

118. Kuo J, Hamid N, Bondar G, Dewing P, Clarkson J, Micevych P. Sex differences in hypothalamic astrocyte response to estradiol stimulation. Biol Sex Differ. 2010;1(1). doi:10.1186/2042-6410-1-7

119. Kuo J, Hamid N, Bondar G, Prossnitz ER, Micevych P. Membrane estrogen receptors stimulate intracellular calcium release and progesterone synthesis in hypothalamic astrocytes. Journal of Neuroscience. 2010;30(39). doi:10.1523/JNEUROSCI.1158-10.2010

120. Beyer C. Nongenomic effects of oestrogen: Embryonic mouse midbrain neurones respond with a rapid release of calcium from intracellular stores. European Journal of Neuroscience. 1998;10(1). doi:10.1046/j.1460-9568.1998.00045.x

121. Ibrahim MMH, Bheemanapally K, Sylvester PW, Briski KP. Sex-specific estrogen regulation of hypothalamic astrocyte estrogen receptor expression and glycogen metabolism in rats. Mol Cell Endocrinol. 2020;504. doi:10.1016/j.mce.2020.110703

122. Pawlak J, Brito V, Küppers E, Beyer C. Regulation of glutamate transporter GLAST and GLT-1 expression in astrocytes by estrogen. Molecular Brain Research. 2005;138(1). doi:10.1016/j.molbrainres.2004.10.043

123. Lee E, Sidoryk-Wêgrzynowicz M, Wang N, et al. GPR30 regulates glutamate transporter GLT-1 expression in rat primary astrocytes. Journal of Biological Chemistry. 2012;287(32). doi:10.1074/jbc.M112.341867

124. Mong JA, Nuñez JL, McCarthy MM. GABA mediates steroid-induced astrocyte differentiation in the neonatal rat Hypothalamus. J Neuroendocrinol. 2002;14(1). doi:10.1046/j.1365-2826.2002.00737.x

125. Spence RD, Wisdom AJ, Cao Y, et al. Estrogen mediates neuroprotection and anti-inflammatory effects during EAE through ERα signaling on astrocytes but not through ERβ signaling on astrocytes or neurons. Journal of Neuroscience. 2013;33(26). doi:10.1523/JNEUROSCI.0886-13.2013

126. Spence RD, Hamby ME, Umeda E, et al. Neuroprotection mediated through estrogen receptor-α in astrocytes. Proc Natl Acad Sci U S A. 2011;108(21). doi:10.1073/pnas.1103833108

127. Yu X, Nagai J, Khakh BS. Improved tools to study astrocytes. Nat Rev Neurosci. 2020;21(3). doi:10.1038/s41583-020-0264-8

128. Haydon PG, Carmignoto G. Astrocyte control of synaptic transmission and neurovascular coupling. Physiol Rev. 2006;86(3). doi:10.1152/physrev.00049.2005

129. Nichols CD, Roth BL. Engineered G-protein coupled receptors are powerful tools to investigate biological processes and behaviors. Front Mol Neurosci. 2013;2(OCT). doi:10.3389/neuro.02.016.2009

130. Xie Y, Wang T, Sun GY, Ding S. Specific disruption of astrocytic Ca2+ signaling pathway in vivo by adeno-associated viral transduction. Neuroscience. 2010;170(4). doi:10.1016/j.neuroscience.2010.08.034

131. Chen N, Sugihara H, Kim J, et al. Direct modulation of GFAP-expressing glia in the arcuate nucleus bi-directionally regulates feeding. Elife. 2016;5(OCTOBER2016). doi:10.7554/eLife.18716

132. Amatruda M, Villavicencio J, Britton G, Horng S. Aldh1l1-cre/ERT2 promoter drives flox-mediated recombination in peripheral immune cells in addition to astrocytes (P2-3.007). Neurology. 2023;100(17_supplement_2). doi:10.1212/wnl.0000000000203599

133. Guttenplan KA, Liddelow SA. Astrocytes and microglia: Models and tools. Journal of Experimental Medicine. 2019;216(1). doi:10.1084/jem.20180200

134. Taschenberger G, Tereshchenko J, Kügler S. A MicroRNA124 Target Sequence Restores Astrocyte Specificity of gfaABC1D-Driven Transgene Expression in AAV-Mediated Gene Transfer. Mol Ther Nucleic Acids. 2017;8. doi:10.1016/j.omtn.2017.03.009

135. Liu Y, Namba T, Liu J, Suzuki R, Shioda S, Seki T. Glial fibrillary acidic protein-expressing neural progenitors give rise to immature neurons via early intermediate progenitors expressing both glial fibrillary acidic protein and neuronal markers in the adult hippocampus. Neuroscience. 2010;166(1). doi:10.1016/j.neuroscience.2009.12.026

136. Su M, Hu H, Lee Y, D’Azzo A, Messing A, Brenner M. Expression specificity of GFAP transgenes. Neurochem Res. 2004;29(11 SPEC. ISS.):2075–2093. doi:10.1007/s11064-004-6881-1

137. Mckinley MJ, Yao ST, Uschakov A, McAllen RM, Rundgren M, Martelli D. The median preoptic nucleus: Front and centre for the regulation of body fluid, sodium, temperature, sleep and cardiovascular homeostasis. Acta Physiologica. 2015;214(1). doi:10.1111/apha.12487

138. Gvilia I, Xu F, McGinty D, Szymusiak R. Homeostatic regulation of sleep: A role for preoptic area neurons. Journal of Neuroscience. 2006;26(37). doi:10.1523/JNEUROSCI.2012-06.2006

139. Kolaj M, Renaud LP. Metabotropic glutamate receptors in median preoptic neurons modulate neuronal excitability and glutamatergic and gabaergic inputs from the subfornical organ. J Neurophysiol. 2010;103(2). doi:10.1152/jn.00808.2009

140. Eyigor O, Centers A, Jennes L. Distribution of ionotropic glutamate receptor subunit mRNAs in the rat hypothalamus. Journal of Comparative Neurology. 2001;434(1). doi:10.1002/cne.1167

141. Zhang J, Yin D, Wu F, et al. Microinjection of adenosine into the hypothalamic ventrolateral preoptic area enhances wakefulness via the A1 receptor in rats. Neurochem Res. 2013;38(8). doi:10.1007/s11064013-1063-7

142. Hayaishi O. Sleep-wake regulation by prostaglandins D2 and E2. Journal of Biological Chemistry. 1988;263(29). doi:10.1016/s0021-9258(18)68073-1

143. Wang F, Smith NA, Xu Q, et al. Astrocytes modulate neural network activity by Ca 2+-dependent uptake of extracellular K +. Sci Signal. 2012;5(218). doi:10.1126/scisignal.2002334

144. Pacholko AG, Wotton CA, Bekar LK. Astrocytes—The Ultimate Effectors of Long-Range Neuromodulatory Networks? Front Cell Neurosci. 2020;14. doi:10.3389/fncel.2020.581075

145. Shah D, Gsell W, Wahis J, et al. Astrocyte calcium dysfunction causes early network hyperactivity in Alzheimer’s disease. Cell Rep. 2022;40(8). doi:10.1016/j.celrep.2022.111280

146. Van Den Herrewegen Y, Sanderson TM, Sahu S, De Bundel D, Bortolotto ZA, Smolders I. Side-by-side comparison of the effects of Gq-and Gi-DREADD-mediated astrocyte modulation on intracellular calcium dynamics and synaptic plasticity in the hippocampal CA1. Mol Brain. 2021;14(1). doi:10.1186/s13041-021-00856-w

147. Chen G, Li HM, Chen YR, Gu XS, Duan S. Decreased estradiol release from astrocytes contributes to the neurodegeneration in a mouse model of Niemann-Pick disease type C. Glia. 2007;55(15). doi:10.1002/glia.20563

148. Hu R, Cai WQ, Wu XG, Yang Z. Astrocyte-derived estrogen enhances synapse formation and synaptic transmission between cultured neonatal rat cortical neurons. Neuroscience. 2007;144(4). doi:10.1016/j.neuroscience.2006.09.056

149. Batiuk MY, Martirosyan A, Wahis J, et al. Identification of region-specific astrocyte subtypes at single cell resolution. Nat Commun. 2020;11(1). doi:10.1038/s41467-019-14198-8

150. Womac AD, Burkeen JF, Neuendorff N, Earnest DJ, Zoran MJ. Circadian rhythms of extracellular ATP accumulation in suprachiasmatic nucleus cells and cultured astrocytes. European Journal of Neuroscience. 2009;30(5):869–876. doi:10.1111/j.1460-9568.2009.06874.x

151. Bojarskaite L, Bjørnstad DM, Pettersen KH, et al. Astrocytic Ca2+ signaling is reduced during sleep and is involved in the regulation of slow wave sleep. Nat Commun. 2020;11(1):3240. doi:10.1038/S41467-020-17062-2

152. Ingiosi AM, Hayworth CR, Harvey DO, et al. A role for astroglial calcium in mammalian sleep and sleep regulation. Curr Biol. 2020;30(22):4373. doi:10.1016/J.CUB.2020.08.052

153. Vo DKH, Hartig R, Weinert S, Haybaeck J, Nass N. G-Protein-Coupled Estrogen Receptor (GPER)-Specific Agonist G1 Induces ER Stress Leading to Cell Death in MCF-7 Cells. Biomolecules. 2019;9(9):503. doi:10.3390/BIOM9090503

154. Revankar CM, Cimino DF, Sklar LA, Arterburn JB, Prossnitz ER. A transmembrane intracellular estrogen receptor mediates rapid cell signaling. Science. 2005;307(5715):1625–1630. doi:10.1126/SCIENCE.1106943

155. Mong JA, Devidze N, Goodwillie A, Pfaff DW. Reduction of lipocalin-type prostaglandin D synthase in the preoptic area of female mice mimics estradiol effects on arousal and sex behavior. Proc Natl Acad Sci U S A. 2003;100(25):15206–15211. doi:10.1073/PNAS.2436540100

156. Poskanzer KE, Yuste R. Astrocytes regulate cortical state switching in vivo. Proc Natl Acad Sci U S A. 2016;113(19):E2675–E2684. doi:10.1073/PNAS.1520759113/-/DCSUPPLEMENTAL

157. Vaidyanathan T V., Collard M, Yokoyama S, Reitman ME, Poskanzer KE. Cortical astrocytes independently regulate sleep depth and duration via separate gpcr pathways. Elife. 2021;10. doi:10.7554/ELIFE.63329

158. Falcone C, Penna E, Hong T, et al. Cortical Interlaminar Astrocytes Are Generated Prenatally, Mature Postnatally, and Express Unique Markers in Human and Nonhuman Primates. Cerebral Cortex. 2021;31(1):379–395. doi:10.1093/CERCOR/BHAA231

159. Degl’Innocenti E, Dell’Anno MT. Human and mouse cortical astrocytes: a comparative view from development to morphological and functional characterization. Front Neuroanat. 2023;17:1130729. doi:10.3389/FNANA.2023.1130729

160. Amateur SK, McCarthy MM. Sexual Differentiation of Astrocyte Morphology in the Developing Rat Preoptic Area. J Neuroendocrinol. 2002;14(11):904–910. doi:10.1046/J.1365-2826.2002.00858.X

161. Bekar LK, He W, Nedergaard M. Locus coeruleus α-adrenergic-mediated activation of cortical astrocytes in vivo. Cerebral Cortex. 2008;18(12):2789–2795. doi:10.1093/cercor/bhn040

162. Ding F, O’Donnell J, Thrane AS, et al. α1-Adrenergic receptors mediate coordinated Ca2+ signaling of cortical astrocytes in awake, behaving mice. Cell Calcium. 2013;54(6):387–394. doi:10.1016/J.CECA.2013.09.001

163. Schnell C, Negm M, Driehaus J, Scheller A, Hülsmann S. Norepinephrine-induced calcium signaling in astrocytes in the respiratory network of the ventrolateral medulla. Respir Physiol Neurobiol. 2016;226:18–23. doi:10.1016/J.RESP.2015.10.008

164. Feinberg I, Maloney T, Campbell IG. Effects of hypnotics on the sleep EEG of healthy young adults: new data and psychopharmacologic implications. J Psychiatr Res. 2000;34(6):423–438. doi:10.1016/S0022-3956(00)00038-8

165. Roehrs T, Roth T. Drug-related Sleep Stage Changes: Functional Significance and Clinical Relevance. Sleep Med Clin. 2010;5(4):559. doi:10.1016/J.JSMC.2010.08.002

166. Guo F, Yi L, Zhang W, Bian ZJ, Zhang YB. Association Between Z Drugs Use and Risk of Cognitive Impairment in Middle-Aged and Older Patients With Chronic Insomnia. Front Hum Neurosci. 2021;15:775144. doi:10.3389/FNHUM.2021.775144

167. González-Reyes RE, Nava-Mesa MO, Vargas-Sánchez K, Ariza-Salamanca D, Mora-Muñoz L. Involvement of astrocytes in Alzheimer’s disease from a neuroinflammatory and oxidative stress perspective. Front Mol Neurosci. 2017;10. doi:10.3389/fnmol.2017.00427

168. Haughey NJ, Mattson MP. Alzheimer’s amyloid β-peptide enhances ATP/Gap junction-mediated calcium-wave propagation in astrocytes. Neuromolecular Med. 2003;3(3). doi:10.1385/NMM:3:3:173

169. Garwood CJ, Ratcliffe LE, Simpson JE, Heath PR, Ince PG, Wharton SB. Review: Astrocytes in Alzheimer’s disease and other age-associated dementias: a supporting player with a central role. Neuropathol Appl Neurobiol. 2017;43(4). doi:10.1111/nan.12338

170. Jiang R, Diaz-Castro B, Looger LL, Khakh BS. Dysfunctional calcium and glutamate signaling in striatal astrocytes from Huntington’s disease model mice. Journal of Neuroscience. 2016;36(12). doi:10.1523/JNEUROSCI.3693-15.2016

171. Park JS, Kam TI, Lee S, et al. Blocking microglial activation of reactive astrocytes is neuroprotective in models of Alzheimer’s disease. Acta Neuropathol Commun. 2021;9(1). doi:10.1186/S40478-021-01180-Z

172. Reiner DJ, Mietlicki-Baase EG, McGrath LE, et al. Astrocytes Regulate GLP-1 Receptor-Mediated Effects on Energy Balance. The Journal of Neuroscience. 2016;36(12):3531. doi:10.1523/JNEUROSCI.3579-15.2016

173. Zheng J, Xie Y, Ren L, et al. GLP-1 improves the supportive ability of astrocytes to neurons by promoting aerobic glycolysis in Alzheimer’s disease. Mol Metab. 2021;47:101180. doi:10.1016/J.MOLMET.2021.101180

174. Batista AF, Forny-Germano L, Clarke JR, et al. The diabetes drug liraglutide reverses cognitive impairment in mice and attenuates insulin receptor and synaptic pathology in a non-human primate model of Alzheimer’s disease. J Pathol. 2018;245(1):85–100. doi:10.1002/PATH.5056

175. Wang ZJ, Han YF, Zhao F, et al. A dual GLP-1 and Gcg receptor agonist rescues spatial memory and synaptic plasticity in APP/PS1 transgenic mice. Horm Behav. 2020;118. doi:10.1016/J.YHBEH.2019.104640

176. Li T, Jiao JJ, Hölscher C, et al. A novel GLP-1/GIP/Gcg triagonist reduces cognitive deficits and pathology in the 3xTg mouse model of Alzheimer’s disease. Hippocampus. 2018;28(5):358–372. doi:10.1002/HIPO.22837

177. Lynch N, Luca R De, Spinieli RL, et al. Identifying the Brain Circuits that Regulate Pain-Induced Sleep Disturbances. bioRxiv. Published online December 20, 2024:2024.12.20.629596. doi:10.1101/2024.12.20.629596

178. Hambrecht-Wiedbusch VS, Gabel M, Liu LJ, Imperial JP, Colmenero A V., Vanini G. Preemptive Caffeine Administration Blocks the Increase in Postoperative Pain Caused by Previous Sleep Loss in the Rat: A Potential Role for Preoptic Adenosine A2A Receptors in Sleep–Pain Interactions. Sleep. 2017;40(9). doi:10.1093/SLEEP/ZSX116

179. Swift KM, Gary NC, Urbanczyk PJ. On the basis of sex and sleep: the influence of the estrous cycle and sex on sleep-wake behavior. Front Neurosci. 2024;18:1426189. doi:10.3389/FNINS.2024.1426189/PDF

180. Scammell TE, Arrigoni E, Lipton JO. Neural Circuitry of Wakefulness and Sleep. Neuron. 2017;93(4). doi:10.1016/j.neuron.2017.01.014

